# Protein complex stoichiometry and expression dynamics of transcription factors modulate stem cell division

**DOI:** 10.1101/439331

**Authors:** Natalie M. Clark, Adam P. Fisher, Barbara Berckmans, Lisa Van den Broeck, Emily C. Nelson, Thomas T. Nguyen, Estefano Bustillo-Avendaño, Sophia G. Zebell, Miguel Moreno-Risueno, Rüdiger Simon, Kimberly L. Gallagher, Rosangela Sozzani

## Abstract

Stem cells divide and differentiate to form all the specialized cell types in a multicellular organism. In the Arabidopsis root, stem cells are maintained in an undifferentiated state by a less mitotically active population of cells called the Quiescent Center (QC). Determining how the QC regulates the surrounding stem cell initials, or what makes the QC fundamentally different from the actively dividing initials, is important for understanding how stem cell divisions are maintained. Here, we gained insight into the differences between the QC and the Cortex Endodermis Initials (CEI) by studying the mobile transcription factor SHORTROOT (SHR) and its binding partner SCARECROW (SCR). We constructed an Ordinary Differential Equation (ODE) model of SHR and SCR in the QC and CEI which incorporated the stoichiometry of the SHR-SCR complex as well as upstream transcriptional regulation of SHR and SCR. Our model prediction coupled with experimental validation showed that high levels of the SHR-SCR complex is associated with more CEI division but less QC division. Further, our model prediction allowed us to establish the timing of QC and CEI division and propose that SHR repression of QC division depends on the formation of SHR homodimer. Thus, our results support that SHR-SCR protein complex stoichiometry and regulation of SHR transcription modulate the division timing of two different specialized cell types in the root stem cell niche.

## Introduction

The tight, coordinated regulation of stem cell division and differentiation ensures proper development and growth of multicellular organisms. In the Arabidopsis root stem cell niche (SCN), controlled asymmetric divisions regenerate the stem cells and produce all of the differentiated tissues from different stem cell populations. The Arabidopsis root SCN also contains a relatively mitotically inactive population of cells, known as the Quiescent Center (QC), which is thought to maintain the surrounding initials in an undifferentiated state, allowing them to undergo asymmetric divisions (1, 2).Thus, determining how the QC regulates the surrounding stem cells, or what makes the QC fundamentally different from the actively dividing initial cells, is important for understanding how stem cell divisions are controlled and maintained during development.

Mobile signals, including transcription factors (TFs), have regulatory roles in the Arabidopsis root SCN (3–5). Specifically, the mobile TF SHORTROOT (SHR) has a key role in controlling the asymmetric division of the Cortex Endodermis Initial (CEI) cells. The CEI cell first divides anticlinally to form the CEI daughter (CEID) cell which then divides periclinally to form the endodermis and cortex layers. Periclinal division can also occur in the CEI itself (6). After SHR is transcribed and translated in the stele, the SHR protein moves to the CEI cells, QC and endodermis where it forms a complex with the transcription factor, SCARECROW (SCR). SHR, and SCR control periclinal division of the CEI/CEID cells (hereafter collectively referred to as the CEI) through the regulation of CYCLIND6;1 (CYCD6;1) (3, 6–13). Members of the C2H2 zinc finger family of TFs (the BIRD proteins) also redundantly regulate this CEI division alongside SCR (9, 14, 15).

Movement of SHR out of the endodermis into the cortex or epidermis is restricted by SCR. In roots with reduced levels of SCR, SHR is able to move past the endodermis and ectopic periclinal division occur in the cortex and epidermis (16–18). Likewise, SHR moves out of the endodermis back to the vasculature in the absence of SCR (19). Therefore, it was suggested that the SHR-SCR complex acts to spatially restrict SHR activity, and thus cell division, to the endodermis. In addition, recent work has shown that two members of the BIRD family named JACKDAW (JKD) and BALDIBIS also bind to SCR to restrict SHR movement past the endodermal cell layer of the ground tissue and confer endodermal fate (14). Thus, the regulation of SHR movement is an important mechanism for restricting cell division in the Arabidopsis root.

SHR also moves into, and binds SCR, in the QC. While recent work used FRET-FLIM to show that both the SHR-SCR and SHR-JKD complexes form in the QC, with the SHR-JKD complex contributing to QC specification and maintenance (9), it is still unclear how the SHR-SCR complex may contribute to QC division as the QC is relatively mitotically inactive (20). One hypothesis is that the oligomeric state of SHR and SCR and stoichiometry of the SHR-SCR protein complex could affect its function, as this is the case with other protein complexes in Arabidopsis (21, 22). It has already been shown that the SHR-SCR stoichiometry may be important for its function in the endodermis, as the SHR-SCR complex exists in both a 1 SHR: 1 SCR and 2 SHR: 1 SCR stoichiometry. Also, it has been shown that SHR homodimer formation in the endodermis depends on the presence of SCR (19). Therefore, determining the oligomeric state and protein complex stoichiometry of SHR and SCR proteins would increase our understanding of their differing roles in QC and CEI division.

Here, we show how differences in expression levels and stoichiometry of the SHR and SCR complex affect their roles in QC and CEI division. We first used scanning FCS to show that SHR and SCR form complexes with 4 different stoichiometries in the QC, as compared to 2 in the CEI. We then incorporated these experimental results into an Ordinary Differential Equation (ODE) model of SHR and SCR expression in the QC and CEI. This model incorporated predicted, upstream regulators of SHR and SCR in addition to SHR and SCR themselves. Using a combination of model simulations and experimental validation, we showed how our model inferred that high levels of SHR-SCR complex promote CEI division but repress QC division, and we were able to determine the relative timing of these divisions. Further, we showed that an increase in QC division is associated with the loss of SHR homodimer. Overall, our model provides a predictive framework for how the cell-specific SHR-SCR complex expression levels and complex stoichiometries regulate the timing of QC and CEI division.

## Results

### Predictive model of SHR and SCR incorporates complex stoichiometry and upstream transcriptional regulation

To address if the temporal expression dynamics, and potentially the biological role, of the SHR-SCR complex differs between the QC and CEI, we developed an Ordinary Differential Equation (ODE) model that predicts SHR and SCR levels in the QC and CEI as well as their oligomeric states. For simplicity, we modeled transcriptional regulation and protein expression in the same equation. Our model consists of 20 equations, 17 of which predict the expression of SHR, SCR, the SHR-SCR complex, and the different oligomeric states/stoichiometries of these proteins/protein complexes in the vasculature, CEI and QC. We assumed that SHR and SCR can oligomerize with themselves to form homodimers, as well as heterodimerize with each other to form protein complexes with various stoichiometries (e.g. 1 SHR: 1 SCR, 2 SHR: 1 SCR, etc.) (19). The remaining three equations predict the expression of a predicted, upstream SHR activator, SHR repressor, and SCR repressor as it has been previously shown that some genes may act upstream of this pathway (22). In addition, we incorporated the known upstream activation of SCR by the SHR: SCR complex (17).

The model incorporates 26 parameters, including the previously determined SHR movement from the stele into the QC and CEI (3, 19). To identify the most important parameters in the model and, thus, the additional parameters that needed to be experimentally determined or estimated from data, we first performed a global sensitivity analysis (see Methods, Supplementary Figure 1, Supplementary Table 1). We found that 14 out of the 26 parameters were sensitive, which include: the SHR diffusion coefficient, the formation of SHR and SCR homodimers and heterodimers, the production rate of SHR, and the production rates of the unknown upstream regulators. We set the remaining 12 parameters to constant values, as variations in these parameter values do not significantly impact the model outcome.

We determined SHR and SCR oligomeric state and complex stoichiometries using Number and Brightness (N&B; see Methods) and quantified the percentage of monomeric and homodimeric forms of either SHR or SCR in roots expressing either SHR:SHR-GFP or SCR:SCR-GFP translational fusions. We found that there were significantly more SHR and SCR homodimers in the QC (16.9% SHR homodimer, 17.3% SCR homodimer) than in the CEI (11.3% SHR homodimer, 6.6% SCR homodimer) (Figure 1A). To determine if the stoichiometry of SHR-SCR complexes varied between the QC and CEI, we performed Cross-Correlation Number & Brightness (Cross N&B; see Methods) on roots expressing both the SHR:SHR-GFP and SCR:SCR-mCherry translational fusions (Figure 1B). The output of Cross N&B is a stoichiometry histogram that gives qualitative information about the amount of SHR-SCR complex formation. To quantitatively assess the different protein complexes, we developed a binding score using this histogram in which a high score (6, red line) corresponded to a high association between SHR and SCR and a low score (1 or 0, dark blue line or no line) corresponded to no association between SHR and SCR (Supplementary Figure 2, see Methods). We first performed Cross N&B on the vasculature tissue, which does not contain SCR, as a control and found that the proportion of SHR bound to SCR is less than 1% (Supplementary Figure 2). We then found that the number of different stoichiometries, as well as their abundance, varies between the QC and the CEI. First, the 1 SHR: 1 SCR complex has the highest binding score and constitutes the majority (over 90%) of the SHR-SCR complex stoichiometries in both the CEI and QC (Figure 1B-C). Second, the complexes incorporating SCR homodimer (namely 1 SHR: 2 SCR and 2 SHR: 2 SCR) had higher binding scores in the QC than the CEI, suggesting that these complexes are more likely to form in the QC. (Figure 1B). In support of this, we found that the second most abundant complex in the QC is 1 SHR: 2 SCR, (∼7% of complexes) in contrast with the 2 SHR: 1 SCR complex (∼7% of complexes) in the CEI. In addition, the 2 SHR: 2 SCR complex does not form in the CEI (binding score ∼1, <1% of complexes) (Figure 1B). Thus, both the percentage of oligomeric states and stoichiometries of the SHR-SCR complex are significantly different between the QC and CEI, suggesting different roles for SHR and SCR in these two cell types.

**Figure 1:**
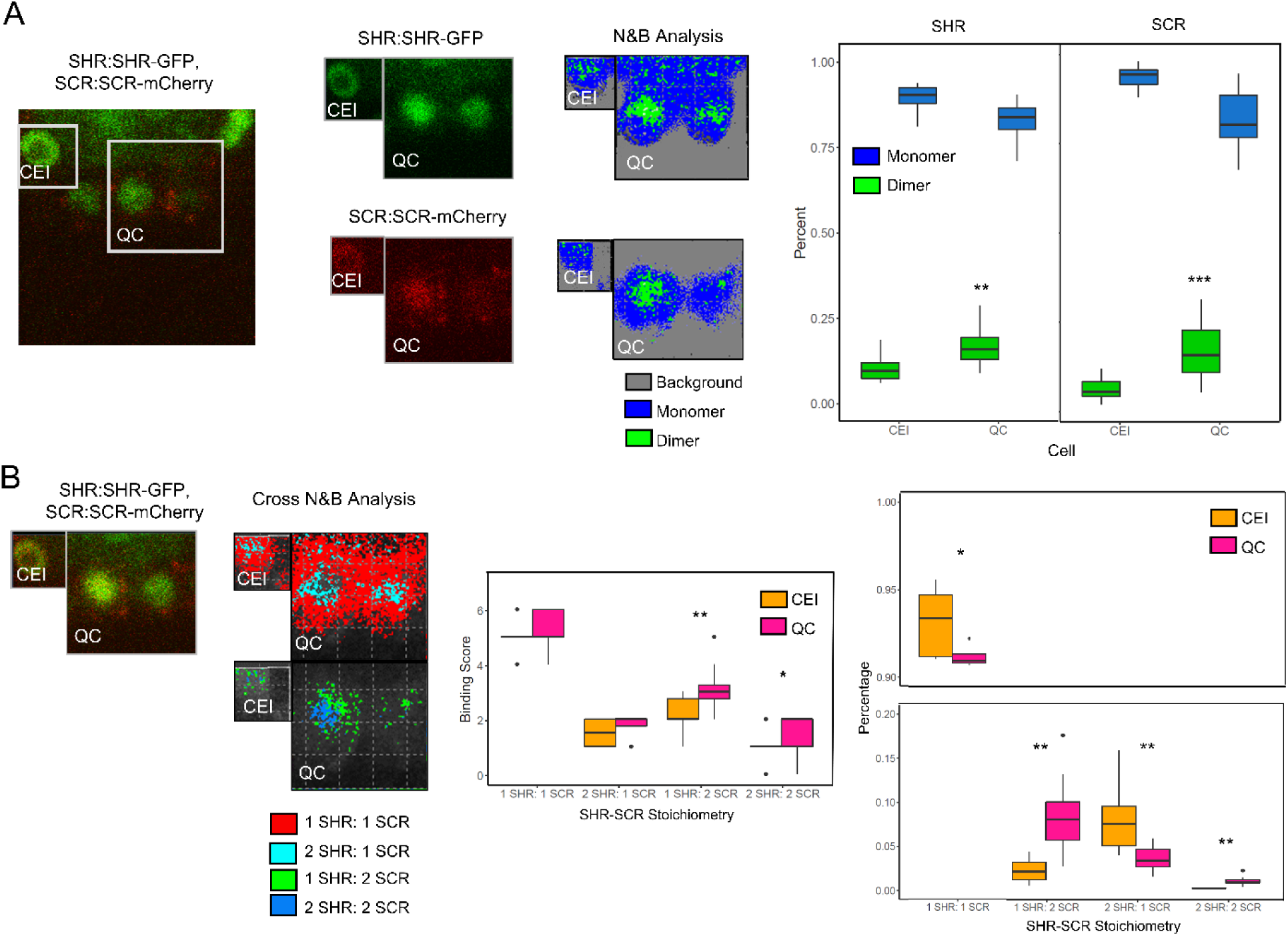
SHR-SCR oligomeric state and stoichiometry differs between CEI and QC. (A) (left) Representative image of SHR:SHR-GFP, SCR:SCR-mCherry in the stem cell niche. Boxes show ROI used for analysis (middle left) SHR:SHR-GFP (top) and SCR:SCR-mCherry (bottom) in the QC (right) and CEI (left). (middle right) N&B analysis of images on the left. Grey represents background fluorescence, blue monomer, and green dimer. (right) Quantification of oligomeric state of SHR (left, n=15) and SCR (right, n=14). ** denotes p < 0.05, *** denotes p < 0.01, Wilcoxon test. (B) (left) Representative image of SHR:SHR-GFP, SCR:SCR-mCherry in the QC (right) and CEI (left). (second from left) Cross N&B analysis of image on the left. Red represents 1 SHR: 1 SCR, light blue represents 2 SHR: 1 SCR, green represents 1 SHR: 2 SCR and dark blue represents 2 SHR: 2 SCR complex. (third from left) Quantification of complex binding scores in the CEI (orange, n=12) and QC (pink, n=12). (right) Quantification of the percentage of SHR-SCR complex stoichiometries in the CEI (orange, n=12) and QC (pink, n=12). There is a break in the graph from 0.2 to 0.9. ** denotes p < 0.05, * denotes p < 0.1, Wilcoxon test.

Given the complete absence of the 2 SHR: 2 SCR complex and the relatively low (<5%) amount of 1 SHR: 2 SCR complex in the CEI, we simplified the model and assumed that these complexes do not form in the CEI. We then estimated the best values for the sensitive parameters. First, we estimated the formation parameters for the SHR and SCR homo- and heterodimers based on our experimentally measured proportions of these complexes (Figure 1B, Supplementary Table 2). Second, we incorporated our previously measured movement coefficient of SHR into the QC and CEI (19). Third, we estimated the production parameters of the upstream SHR activator, SHR repressor, and SCR repressor by fitting our model to SHR and SCR expression obtained from a time course of a stem cell-enriched population in 4 day to 6 day old plants (hereinafter referred to as the stem cell time course) (Supplementary Table 3). Briefly, we collected a population of predominantly root stem cell initials every 8 hours from 4 days to 6 days post-stratification (23). We found that this model incorporating all these aspects of SHR and SCR regulation accurately recapitulated our expression data (Supplementary Figure 2), allowing us to predict how SHR and SCR expression varies temporally in the QC and CEI.

### Model prediction identifies putative upstream regulators of SHR

Our model predicted both the expression of SHR and SCR and their putative upstream regulators (Supplementary Figure 3). Therefore, we leveraged the model prediction and experimental data to infer the most suitable candidates for the upstream SHR regulators. First, we generated a list of 21 transcription factors (TFs) that were either shown to directly bind the SHR promoter (through Chromatin Immunoprecipitation (ChIP) or Yeast-1-Hybrid (Y1H), or whose loss-of-function/overexpression lines show a decrease/increase in SHR expression (23–26) (Supplementary Figure 4, Supplementary Table 4). We then compared the temporal expression of these TFs from our stem cell time course to the model prediction for the SHR activator. To quantify the goodness of fit between the experimental expression and model prediction, we calculated a sign score for each TF, which measures whether the experimental data and the model prediction vary in the same direction. A sign score of 1 corresponded to when the model and expression of the TF changed in the same direction (either increasing or decreasing); whereas a sign score of −1 corresponded to a gene whose expression changed in the opposite direction predicted by the model (see Methods). As a result, we identified 6 genes, STOREKEEPER 1 (STK01), HOMEOBOX 13 (HB13), SEUSS (SEU), bZIP17, ETHYLENE RESPONSIVE ELEMENT BINDING FACTOR 4 (ERF4), and AT3G60580, which all had the highest possible sign score (sign score = 5) and represented candidates for activators of SHR (Supplementary Figure 4, Supplementary Table 4).

Given the role of SHR and SCR in stem cell function, we reasoned that our putative SHR activators should also be expressed in the stem cell niche. Thus, using a stem-cell-type-specific transcriptional dataset, we checked their expression in the xylem initials, as these positive regulators must be expressed in the vasculature in order to transcriptionally regulate SHR (27). Accordingly, we removed HB13 and AT3G60580 from our list given their low expression in the xylem initials (Figure 2A). We additionally removed ERF4 given its annotated role as a transcriptional repressor (28), making it a possible false positive for the SHR activator. For the remaining 3 putative activators, bZIP17, STK01, and SEU, we determined whether SHR expression is affected in their loss-of-function mutant lines. Previous work showed that *bzip17* and *stk01* have an approximately 2-fold decrease in levels of SHR as measured by qPCR (23). We also performed RNA-seq in the *seu-3* mutant and identified SHR among the differentially regulated genes in the mutant background compared to WT (Figure 2B, Supplementary Table 5). We next examined if these 3 mutants showed a stem cell niche phenotype, as this would suggest these TFs specifically regulate SHR in the stem cells. While neither *stk01* nor the *bzip17* mutants showed phenotypes, we found that *seu-3* mutants showed improperly maintained columella stem cells (as shown by starch granules accumulation in columella layer just below the QC), a disorganized stem cell niche, more QC divisions, and very few CEI that have not undergone periclinal divisions (hereinafter referred to as undivided CEI) (Figure 2B, Supplementary Figure 4). Further, it has been shown that SEU directly binds the SHR promoter (26). Therefore, we identified SEU as the putative SHR activator in our model.

**Figure 2:**
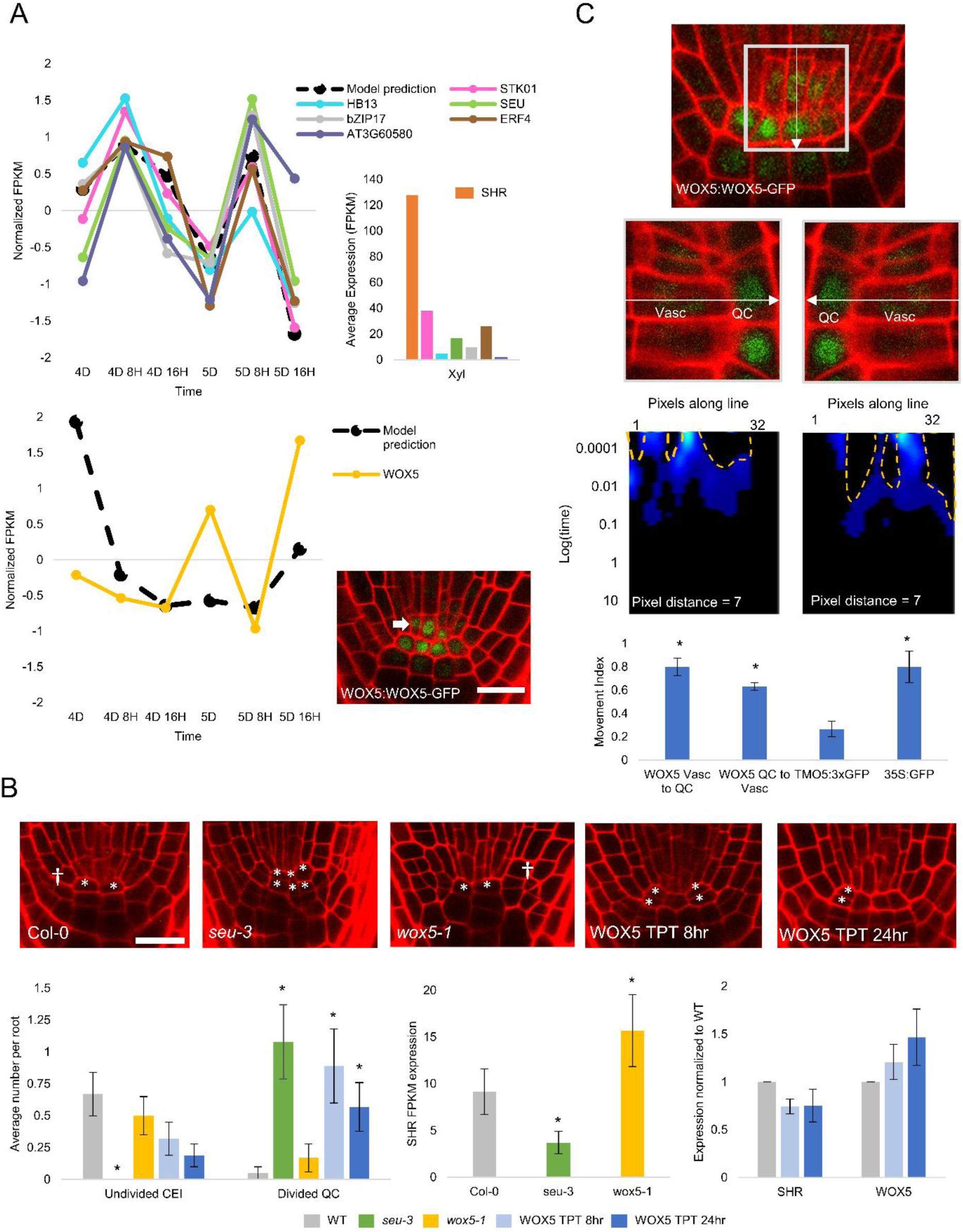
Model identifies putative SHR regulators. (A) (left) Black, dashed line indicates the normalized FPKM value of a SHR activator (top) and SHR repressor (bottom) as predicted by the model using the time course dataset. Colored lines represent the normalized FPKM value of the genes with the maximum possible sign score (i.e. best candidates). (top right) FPKM value of candidate SHR activators in the xylem initials (Xyl) compared to SHR (orange). (bottom right) WOX5:WOX5-GFP expression in the vascular initials (white arrow). Scale bar = 20 µm. (B) (top) Confocal images, from left to right: wild-type root stem cell niche, *seu-3* mutant, *wox5-*1 mutant, WOX5 TPT line after 8 hour induction with beta-estradiol, WOX5 TPT line after 24 hour induction with beta-estradiol. * denotes QC cells, dagger indicates undivided CEI. Scale bar = 20 µm. (bottom left) Quantification of undivided CEI and divided QC in genotypes shown in the top panel. * denotes p<0.05, Wilcoxon with Steel Dwass using Col-0 as control. bottom(bottom middle) SHR expression in *seu-3* and *wox5-*1 mutants. * denotes q < 0.05 and fold change > 2 vs Col-0 (Cuffdiff). Error bars represent 95% confidence intervals returned by Cuffdiff. (bottom right) qPCR of SHR and WOX5 expression in the TPT line 8 (light blue) and 24 (dark blue) hours after induction with beta-estradiol. Error bars represent standard deviation. (C) (top) Image of WOX5:WOX5-GFP root showing the location and direction of the line scan. (second from top) Representative image of WOX5:WOX5:GFP line used for pCF with Quiescent Center (QC) and Vasculature (Vasc) marked. White line shows direction of movement tested. (third from top) pCF carpet of the top panel images. Orange, dashed regions represent arches in the pCF carpet, which denote movement. (bottom) Movement index of WOX5:WOX5:GFP, an immobile control (TMO5:GFP), and a mobile control (35S:GFP) between the QC and vasculature. * denotes p<0.05 as compared to non-mobile control using Wilcoxon with Steel Dwass. Error bars are SEM.

We next focused on identifying putative repressors of SHR expression using the sign score measurement and obtained a list of 25 candidates from similar sources as the SHR activators (Supplementary Table 4). Out of these 25 putative repressors, only NAC13 had the highest possible sign score of 5. However, NAC13 expression is enriched in the mature xylem cells compared to the meristematic vasculature cells (Supplementary Figure 5) (29). This suggests that NAC13 likely does not repress SHR in the stem cell niche region. Thus, to identify repressors of SHR in the stem cell niche, we searched for genes expressed broadly in the stem cell region. To do this, we obtained a list of 125 TFs which are significantly enriched in the stem cells compared to a population of non-stem cells (27) and calculated their sign score when compared to the model prediction of the SHR repressor (Supplementary Table 6, Supplementary Figure 5). WUSCHEL-RELATED HOMEOBOX 5 (WOX5) was the only TF that had the highest possible sign score of 5 (Figure 2B, Supplementary Table 6), suggesting that WOX5 is a putative repressor of SHR.

While WOX5 transcript is largely absent in the stele (28–30), a WOX5:WOX5-GFP translational fusion shows WOX5 protein expression in the vasculature initials (31) (Figure 2B). As WOX5 has been shown to be a mobile protein (30), we postulated that WOX5 protein localization in the vascular initials could be due to WOX5 movement from the QC (Figure 2C). To this end, we performed Pair Correlation Function (pCF) analysis on the WOX5:WOX5-GFP translational fusion and compared its movement to a protein that is able to move (35S:GFP) and a protein that is not able to move (TMO5:3xGFP) from the QC to the vascular initials (19) (see Methods). We found that the WOX5 protein moves from the QC to the vascular initials with a movement index (MI) of 0.75 ± 0.06, which is significantly higher than the movement index of our non-mobile control protein (TMO5:3xGFP, MI= 0.27 ± 0.07) (Figure 2C). These data are consistent with WOX5 movement from the QC to the vascular initials.

To test if WOX5 is a putative regulator of SHR, we performed RNA-seq on a *wox5-1* mutant line and found that SHR is one of the 3302 genes that had higher expression when compared to wild type, suggesting that WOX5 represses SHR (Figure 2B, Supplementary Table 7). Additionally, we obtained a TRANSPLANTA (TPT) overexpression line (32) of WOX5 and observed reduced SHR levels 8 and 24 hours after induction of WOX5 overexpression with beta-estradiol (Figure 2B, see Methods). We quantified both QC and CEI divisions in these two lines as we did for the *seu-3* mutant. While we did not see any significant change in QC or CEI divisions in the *wox5-1* mutant, we did find more QC divisions and less undivided CEI in the WOX5 TPT line (Figure 2B). Therefore, we considered WOX5 as the putative SHR repressor in our model.

### SHR-SCR complex levels modulate the timing of CEI division

After updating the parameters in our model using SEU and WOX5 expression data (Supplementary Table 8, Supplementary Figure 6), we examined our model prediction of SHR and SCR levels in the CEI. Our model predicts that both stoichiometries of the SHR:SCR complex reach their first maximum value between 4 days 8 hours (4D 8H) and 4 days 16 hours after stratification (4D 16H) (Figure 3A, Supplementary Figure 6). Based on previous results showing that high levels of the SHR-SCR complex trigger CEI division, and that CEI divisions occur every 18-24 hours (6), we hypothesized that high levels of the SHR-SCR complex induce CEI division between 4D 16H and 5D. To test this hypothesis, we observed the expression of the CYCD6 marker (pCYCD6::GUS-GFP), which is expressed immediately preceding CEI division (7), in 5D old roots and found that 77.8% of roots show CYCD6:GUS-GFP expression, suggesting that most of the CEI are undergoing periclinal division at 5D (Figure 3C). Between 5D and 6D, the pattern of SHR:SCR complex expression is similar, except that the total complex level is predicted to be 1.5-fold lower between 5D 16H and 6D compared to 4D 16H and 5D. To validate the model prediction, we quantified the fluorescence of the SHR:SHR-GFP and SCR:SCR-GFP markers in the CEI and found that there is less SHR and SCR fluorescence after 5D 16H (Figure 3B). We reasoned that the relatively low levels of the SHR-SCR complex during this time would correlate with less actively dividing CEI. Accordingly, we observed that only 18.2% of 6D roots expressed the CYCD6 marker compared to 77.8% of the roots at 5D (Figure 3C). Further, this decrease in CYCD6 expression is associated with less undivided CEI (27.8% at 5D 16H vs 10.4% at 6D 16H), suggesting that most of the CEI have undergone their periclinal divisions and are no longer actively dividing (Figure 3C). Altogether, our model and experimental data support that the SHR:SCR regulated periclinal division of the CEI halts around 6D.

**Figure 3:**
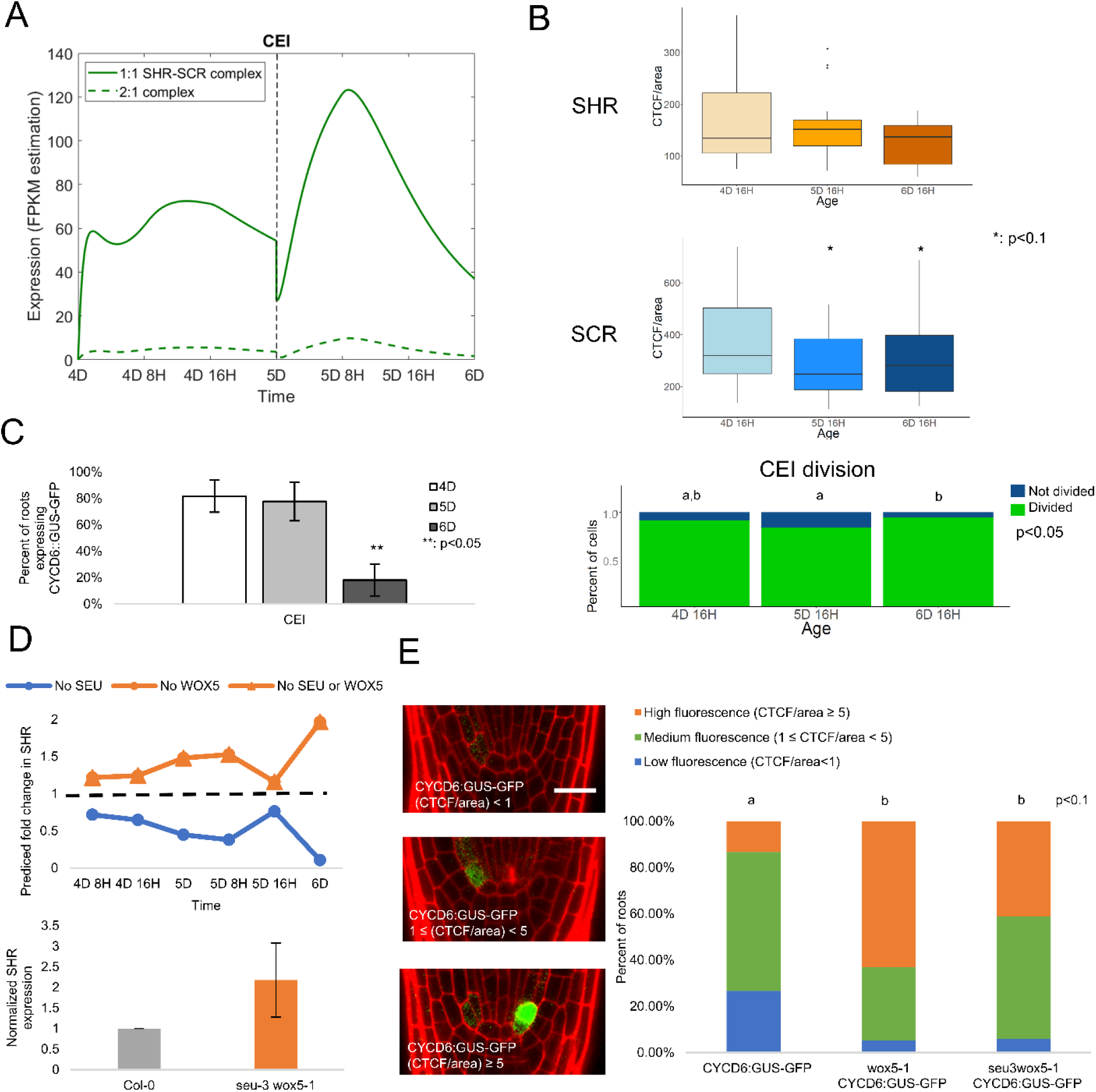
SHR and SCR temporal dynamics modulate CEI division. (A) Model simulation of the expression of SHR/SCR complexes in the endodermis. Black dashed line demarcates CEI division at 5D. Green, solid line represents 1 SHR: 1 SCR complex. Green, dashed line represents 2 SHR: 1 SCR complex. (B) Corrected Total Cell Fluorescence (CTCF) of SHR:SHR-GFP (top) and SCR:SCR-GFP (bottom) in the CEI in 4 day 16 hour (4D 16H), 5 day 16 hour (5D 16H), and 6 day 16 hour (6D 16H) roots. * denotes p<0.1, Wilcoxon with Steel Dwass using 4D 16H as a control. (C) (left) Observed changes in the percent of roots expressing pCYCD6::GUS-GFP. ** denotes p < 0.05, Wilcoxon with Steel Dwass using 4D 16H as a control. Error bars are SEM. (right) Quantification of the percent of periclinally divided CEI (green) in 4D 16H, 5D 16H, and 6D 16H roots. Different letters represent groups that are statistically significantly different at the p<0.05 level using a Chi Squared test with likelihood ratio. The no WOX5 and no SEU or no WOX5 curves overlap. (D) (top) Model prediction of fold change in *SHR* expression in different mutant backgrounds. Blue line represents no SEU production. Orange line represents no WOX5 production. Orange line with triangles represents no SEU or WOX5 production. Black, dashed line indicates a fold-change of 1, where the expression would be equal to the wild-type model prediction. (bottom) SHR expression in *seu-3 wox5-1* mutants measured using qPCR. Results are from two biological replicates. (E) (left) Representative images of CYCD6::GUS-GFP with low fluorescence (CTCF/area) < 1, medium fluorescence 1 < (CTCF/area) < 5, high fluorescence (CTCF/area) > 5 (right half). Scale bar = 20 µm. (right) Quantification of CYCD6:GUS-GFP expression in wild-type (Col-0, n=15), *wox5-1* (n=19), and *seu-3;wox5-1* (n=17) lines. Blue part of bar represents low fluorescence. Different letters represent groups that are statistically significantly different at the p<0.05 level using a Chi Squared test with likelihood ratio.

We also observed a decrease in the number of undivided CEI in both the *seu-3* mutant and WOX5 TPT overexpression lines, which have lower SHR levels (Figure 2B). Thus, we used our model to investigate the effect of SEU and WOX5 on SHR dynamics and consequently CEI division. We first simulated how SHR expression changes in the model when we individually removed SEU or WOX5 production. The model predicted that roots with no SEU production should have lower SHR expression, while roots with no WOX5 production should have more SHR expression (Figure 3D). This agreed with our biological data, which show lower SHR expression in *seu-3* mutants and higher SHR expression in *wox5-1* mutants (Figure 2B). We next used the model to simulate SHR expression upon removing both SEU and WOX5 production as this would be representative of a double mutant. Our model suggested that SHR expression increased in the absence of both SEU and WOX5 to the same extent as when there is no production of only WOX5, such that SHR expression in *seu-3 wox5-1* double mutants should be the same as in *wox5-1* single mutants (Figure 3D). To test the accuracy of the model prediction, we performed qPCR on the *seu-3 wox5-1* double mutant and compared the change in SHR to our RNA-seq data of the *wox5-1* single mutant. We found that *seu-3 wox5-1* mutants have approximately 2-fold higher SHR expression (Figure 3D), which was a similar increase in SHR compared to the *wox5-1* single mutant (1.7-fold, Figure 2B).

Finally, we examined the behavior of our CYCD6 marker in these mutants to infer how these changes in SHR expression affect CEI division. At 5D, we found that *wox5-1* mutants had more roots with high CYCD6 expression compared to wild-type plants at 5D (63.2% of *wox5-1* mutant roots compared to 13.3% of wild-type roots) (Figure 3E), which supports our model prediction that less WOX5 production results in a higher SHR expression and more actively dividing CEI. While *seu-3 wox5-1* double mutants also had a higher percentage of roots with high CYCD6 expression compared to wild-type (41.2% of *seu-3 wox5-1* roots), this proportion of roots with high expression was not statistically different from the *wox5-1* single mutant roots (Figure 3E), which supports the additional model prediction that the double mutant has the same effect as the single *wox5-1* mutant. Overall, these results suggest that WOX5 and SEU coordinate together to control the levels of SHR expression in the CEI.

### Loss of SHR homodimer is associated with QC division

We next found that, in the QC cells, our model predicts the levels of the SHR-SCR complex greatly decrease between 5D 16H and 6D (Figure 4A). As in the CEI, we were able to validate this predicted decrease in SHR and SCR by examining SHR:SHR-GFP and SCR:SCR-GFP expression in the QC cells (Figure 4B). We also found that at the developmental time point that the SHR:SCR complex drastically decreased (5D 16H), more QC divisions were observed compared to an earlier time point in development (4D 16H) (Figure 4B). To further strengthen the connection between the abundance of the SHR-SCR complex and QC divisions, we treated SHR:SHR-GFP and SCR:SCR-GFP lines with brassinolide (BL) as this has been shown to induce QC divisions (31) (Supplementary Figure 7). We found that this BL-induced increase in QC divisions was accompanied by significantly less SHR:SHR-GFP and SCR:SCR-GFP fluoresence in the QC (Supplementary Figure 7). Thus, a decrease in SHR-SCR complex expression promotes QC division, which contrasts with its role in the CEI.

**Figure 4:**
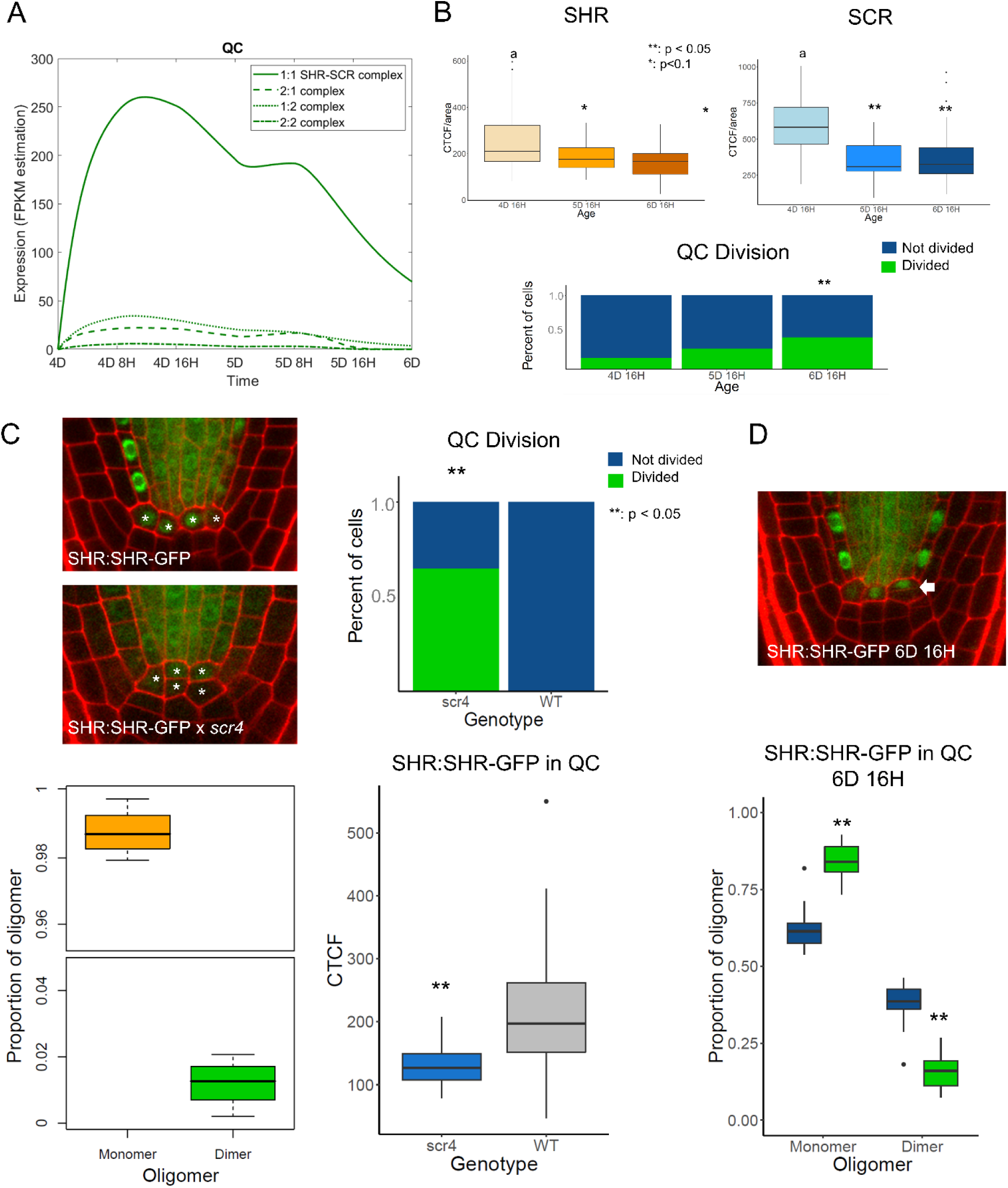
Less SHR expression and loss of SHR homodimer correlate with QC division. (A) Model simulation of the expression of SHR/SCR complexes in the QC. Green solid line represents 1 SHR: 1 SCR. Green dashed line represents 2 SHR: 1 SCR. Green dotted line represents 1 SHR: 2 SCR. Green dot/dashed line represents 2 SHR: 1 SCR. (B) (top) Corrected Total Cell Fluorescence (CTCF) of SHR:SHR-GFP (left) and SCR:SCR-GFP (right) in the QC in 4 day 16 hour (4D 16H), 5 day 16 hour (5D 16H), and 6 day 16 hour (6D 16H) roots. * denotes p<0.1, ** denotes p<0.05 Wilcoxon with Steel Dwass using 4D 16H as a control. (bottom) Quantification of the percent of divided QC (green) in 4D 16H, 5D 16H, and 6D 16H roots. ** denotes p<0.05,Chi Squared test with likelihood ratio using 4D 16H as a control. (C) (top left) Representative images of SHR:SHR-GFP (top) and SHR:SHR-GFP x *scr4* (bottom) at 5D 16H. * denotes QC cells. (top right) Quantification of the percent of divided QC (green) in *scr4* and WT roots. ** denotes p<0.05, Chi Squared test with likelihood ratio. (bottom left) N&B analysis of SHR:SHR-GFP in *scr4*. Orange represents monomer, green represents dimer. There is a break in the graph between 0.05 and 0.95. (bottom right) CTCF of SHR:SHR-GFP in the QC in *scr4* (blue) and WT (gray) background. ** denotes p<0.05, Wilcoxon test. (D) (top) Representative images of SHR:SHR-GFP at 6D 16H. White arrow denotes a divided QC. (bottom) N&B analysis of SHR:SHR-GFP in divided (green) and undivided (navy) QC cells at 6D 16H. ** denotes p<0.05, Wilcoxon test..

Another difference between the model prediction in the QC and CEI was the proportion of SHR:SCR complex stoichiometries. Our model predicted that the 1 SHR: 1 SCR and 2 SHR: 1 SCR stoichiometries are always expressed (expression > 0) in the CEI, even at 6D when the CEI divides less frequently (Figure 3A, Supplementary Figure 6). In contrast, in the QC, our model predicted that the two complexes incorporating SHR homodimer, namely 2 SHR:1 SCR and 2 SHR: 2 SCR, are not present (expression = 0) at 6D (Figure 4A, Supplementary Figure 6). Since treating the roots with BL increased the number of QC divisions, we performed Cross N&B on SHR and SCR in BL-treated roots to determine whether the relative levels of the different SHR-SCR complex stoichiometries are altered when there are more QC divisions. As the model predicted, we found both the 2 SHR:1 SCR and 2 SHR: 2 SCR complex no longer formed in the QC of BL-treated plants (Supplementary Figure 7). Further, we examined SHR:SHR-GFP plants and found that only a relatively small amount of SHR homodimer (< 2%) forms in BL-treated plants, suggesting that the loss of these stoichiometries in BL-treated plants may be due to the loss of SHR homodimer (Supplementary Figure 7). Therefore, our results suggest that repression of QC division may depend on the SHR-SCR complex stoichiometries that incorporate the SHR homodimer, namely the 2 SHR: 1 SCR and 2 SHR: 2 SCR complexes.

We previously found that the formation of the SHR homodimer in the endodermis depends on the presence of SCR by showing that less than 2% SHR homodimer forms in the endodermis of the SCR RNAi (SCRi) line (19). Accordingly, we performed N&B on SHR:SHR-GFP in the *scr4* mutant (see Methods) and found that there is less than 1% of SHR homodimer in the QC (Figure 4C). In addition, we found that this mutant has less SHR:SHR-GFP expression in the QC and more QC divisions (Figure 4C). Finally, we performed N&B on SHR:SHR-GFP in divided versus undivided QC in wild type conditions where SHR homodimer can form normally. To do this, we used plants that were 6D 16H old, as these have the highest number of QC divisions (Figure 4B). We found that there is significantly more SHR homodimer in undivided QC (36.8% homodimer) than in divided QC (16.0% homodimer) (Figure 2D). Along with our results using BL treatment, this supports that lower SHR expression, less SHR homodimer formation are associated with more QC division. Additionally, this supports that the SHR-SCR complex can repress QC division, since this complex cannot form in the *scr4* mutant.

## Discussion

SHORTROOT (SHR) and SCARECROW (SCR) are known regulators of the asymmetric cell divisions of the Cortex Endodermis Initials (CEI), and while some of their roles in QC specification have been described (32), their role in division of the less mitotically active cells of the Quiescent Center (QC) is less understood. By combining mathematical modeling and experimental data, we showed how both the levels of SHR and SCR transcripts and differences in the stoichiometry of the SHR-SCR complex are associated with differences in CEI and QC division. Together, our model and data support that high levels of the SHR and SCR complex promote CEI but repress QC divisions (Figure 5).

**Figure 5:**
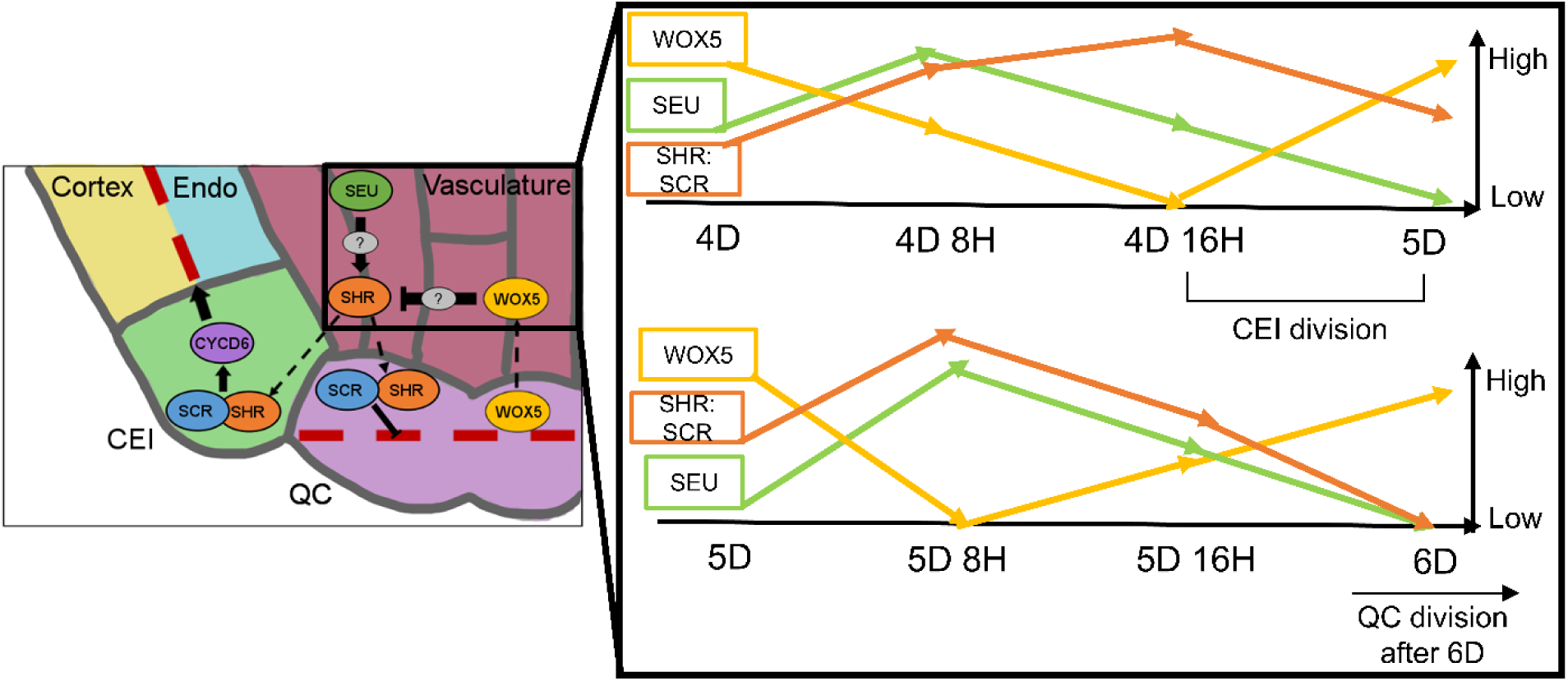
A SEU-WOX5-SHR network modulates the timing of CEI and QC division. (Left) Schematic of the Arabidopsis root stem cell niche. SHR protein (orange) moves from the vasculature into the CEI and QC, where it binds SCR. In the CEI, SHR::SCR activates CYCD6 to promote the periclinal division of the CEI daughter (red, dashed line). In the QC, SHR::SCR represses QC division (red, dashed line). WOX5 protein (yellow) moves from the QC to the vasculature to repress SHR, and SEU protein (green) activates SHR in the vasculature. Both of these regulators may act directly on SHR or indirectly through unknown intermediates (gray). Right(Right, inset) Protein dynamics in the vasculature. The combined activation and repression of WOX5 and SEU, respectively, causes temporal changes in SHR:SCR complex expression. High SHR:SCR complex expression between 4D 16H and 5D coordinates the periclinal division of the CEI daughter. Low SHR:SCR complex expression at 6D correlates with QC division after 6D.

Sensitivity analysis on our mathematical model identified several parameters that were important for predicting SHR and SCR expression dynamics in the CEI and QC, specifically the production and degradation rates of the different oligomeric states and protein complex stoichiometries. Other examples of proteins that form higher oligomers to successfully exert their biological function are Arabidopsis ethylene receptors, which use disulfide bonds to form higher oligomeric states. Disrupting these disulfide bonds greatly affects the receptors’ ability to bind ethylene (22). In another case, the receptors CLAVATA 1 (CLV1) and ARABIDOPSIS CRINKLY 4 (ACR4) form homo- and heterodimeric complexes which differ in their predicted stoichiometries depending on their localization at the plasma membrane or plasmodesmata (33). Using the SHR:SHR-GFP translational fusion in the *scr4* mutant background, we show that the formation of SHR homodimers in the CEI depends on the presence of SCR, which agrees with our previous results in the endodermis (19). Future work could investigate if the binding of SCR to SHR causes a conformational change in SHR that allows it to form a homodimer, contributing to cell-type specific functions.

In addition to using our model to determine how SHR and SCR regulate CEI and QC division, we were also able to predict the timing of these divisions. Our model predicts that low levels of SHR and SCR correlate with QC division (Figure 4). While previous work has shown that SCR has a role in QC identity (34), this is a new insight on how these two genes may regulate QC division. In addition, previous work has shown that the QC cells divide at roughly half the rate of the CEI (20). Therefore, given its slower division rate, we can only speculate that the QC takes longer to divide, perhaps dividing at up to 7 days old (Figure 5). While we have a marker of CEI division (CYCD6), we currently do not have a similar marker for the QC. Moreover, previous work has shown that brassinolide (BL) treated plants have excessive QC divisions that correlate with a lower level of the cell cycle inhibitor ICK2 (35). Given our results showing that BL-treated plants also have lower SHR and SCR levels (Figure 4B), it would be important but beyond the scope of this work to investigate whether SHR and SCR regulate ICK2 to inhibit division of the QC. Additional work could be done to study the role of CYCLIN D 3;3 (CYCD3;3), which is normally expressed outside of the QC but is expressed in the QC in a *wox5-1* mutant (36). We propose that WOX5 represses SHR in the vascular initials, so it is possible that the increase in CYCD3;3 in *wox5-1* mutants could be due to an increase in SHR levels. Finally, mutants in CELL CYCLE SWITCH 52 A2 (CCS52A2) have excessive QC divisions and a disorganized stem cell niche(37, 38), so SHR and SCR could additionally activate CCS52A2 to prevent QC division. Thus, future work could identify whether SHR and SCR regulate ICK2, CYCD3;3, CCS52A2, or other cell cycle genes in the QC to control its division.

We were able to leverage our model to identify two putative SHR regulators, namely SEUSS (SEU) and WUSCHEL-RELATED HOMEOBOX 5 (WOX5). We propose that SEU transcriptionally activates SHR in the vasculature. Additionally, we propose that WOX5 protein moves from the QC into the vascular initials to repress SHR (Figure 5). While it has been shown that SEU can directly bind the SHR promoter (26), it remains unknown whether WOX5 regulation of SHR is direct or indirect. Further, our list of putative SHR regulators consisted of a small, curated list based on previous work. Thus, we do not conclude here that SEU and WOX5 are the only upstream regulators of SHR, and we propose that it is likely that other proteins act in concert with or upstream of SEU and WOX5 to regulate SHR expression in the vasculature. For example, it has been shown that treatment with CLAVATA3/ESR-RELATED 40 peptide (CLE40p) rapidly increases WOX5 protein expression in the vascular initials, and that this CLE40p-dependent upregulation of WOX5 protein expression is also dependent on CLAVATA2 (CLV2) (30). This suggests that there is a gene regulatory network upstream of WOX5 that controls its protein localization, and therefore regulation of SHR, in the vasculature. In addition, SEU is a known transcriptional adapter (39) and therefore likely requires a binding partner to transcriptionally regulate SHR. To identify potential SEU interactors, we completed a Yeast Two-Hybrid (Y2H) screen using SEU as the bait against a TF library generated in (23) (Supplementary Table 9). While we did not exploit these data, because we feel they are outside the scope of this work, we believe it would be important to identify potential binding partners of SEU which are also upstream SHR activators. Finally, SHR is expressed in a gradient, with its highest expression at the tip of the root (11). Thus, potential regulators of SHR may be expressed in the same gradient as SHR, and datasets measuring gene expression gradients in the root could be used to identify additional putative SHR regulators (40). Similar techniques could be used to identify upstream SCR regulators, increasing our understanding of how this expanded SEU-WOX5-SHR-SCR regulatory network controls CEI and QC divisions.

Here, we used a quantitative approach of iterative mathematical modeling with heterogenous experimental methods to show the differing roles of SHR and SCR in the QC and CEI. Specifically, we integrated imaging and gene expression data to quantify SHR-SCR complex stoichiometry, identify putative upstream regulators of SHR, and predict the timing of QC and CEI divisions (Figure 5). While we anticipate that other factors are needed to regulate stem cell division in the Arabidopsis root, our work provides new insights into how SHR and SCR are involved in this process. In conclusion, our results highlight how temporal expression dynamics and changes in protein-protein complex stoichiometry contribute to differential regulation of division in specialized cell types.

## Methods

### Mathematical model formulation and simulation

A system of 21 ordinary differential equations (ODEs) was developed to model the dynamics of SHR and SCR in the CEI and QC (see Supplementary Information for equations). Transcriptional regulation is assumed to happen on a fast time scale such that transcriptional and protein dynamics can be modeled in the same equation. Transcriptional regulation is modeled using Hill equation dynamics, and SHR-SCR complex formation is modeled using mass-action kinetics. SHR diffusion is modeled using a linear term for gradient-independent diffusion. All proteins are assumed to have a linear degradation term. All 4 stoichiometries of the SHR-SCR complex are assumed to activate SCR (16, 19), and it is also assumed that SCR can activate itself (8, 16).

The second version of the model incorporates a Hill equation modeling the regulation of SHR by its activator and repressor. It is assumed that the regulation operates in an “OR” gate such that if the activator and repressor both bind the SHR promoter, the activator overcomes the repressor. This is based on results showing that SHR activator mutants have root phenotypes while SHR repressor mutants do not, suggesting that the activator is stronger than the repressor (23). Another Hill equation models the regulation of SCR by a repressor and the SHR-SCR complex. Finally, the production terms for the SHR activator, SHR repressor, and SCR repressor are assumed to be time-dependent as this produces the best model fit to the experimental data. MATLAB code containing the ODE model is included on the data repository (see Data Availability).

Given the experimental results that the CEI divides at 5D, but the QC does not divide at 5D (Figures 3 & 4), the model was simulated for 4D-5D and 5D-6D separately. To simulate division of the CEI, model values for all of the proteins present in the CEI at 5D were divided by 2. These halved values were then used as the initial values for 5D-6D. It was also assumed that the vasculature divides at 5D, so the same process was repeated for proteins in the vasculature. Since the QC does not divide, protein values in the QC at 5D were used as the initial values for 5D-6D. The model simulations from 4D-5D and 5D-6D were then combined to form the final simulations seen in Figures 3 and 4 and Supplementary Figure 6. MATLAB code used to run these simulations is included on the data repository (see Data Availability).

### Sensitivity analysis of mathematical model

Sensitivity analysis was performed on a version of the model that does not incorporate the QC as the equations for all of the proteins in the CEI and QC are the same: therefore, the sensitivity index of the parameters will be the same in both cell types. The total Sobol effect index (19, 41, 42) was calculated for each parameter value. Parameter values were randomly sampled using Monte Carlo sampling to obtain 150 different values for each parameter (Supplementary Table 10). This process was repeated for 10 technical replicates. The sample number was chosen as 150 as this makes the runtime of the sensitivity analysis approximately 2 hours per technical replicate on an Intel i7-4800MQ 2.70 GHz CPU with 8GB RAM. MATLAB code for calculating the total Sobol index is included on the data repository (see Data Availability). The sensitivity of each variable to each parameter was normalized to [0,1] and then averaged to calculate the final sensitivity indices. The sensitive parameters were chosen as the parameters that had significantly higher Sobol indices than the selected cutoff parameter (*k*_*6*_) using the Wilcoxon test with Steel-Dwass for multiple comparisons with a significance cutoff of p<0.05.

### Parameter estimation

The initial values for *S*_*v*_ (SHR monomer in vasculature), *C*_*e*_ (SCR monomer in CEI), and *C*_*q*_ (SCR monomer in QC) were determined using the FPKM values for SHR and SCR at 4D from replicate 1 of the time course dataset. Since SCR is expressed in both the CEI and QC, the FPKM value needed to be split between the two cells. To accomplish this, expression of SCR was obtained from a dataset of Arabidopsis root tissues (29). It was determined that 41.91% of SCR is expressed in the QC, while the remaining 58.09% is expressed in the CEI and endodermis (Supplementary Table 10). In addition, the results from the N&B analysis (Figure 1) were used to set the initial value of *C*_*2q*_ (SCR dimer in the QC). Once SEU and WOX5 were incorporated into the model, their FPKM values at 4D were used for the initial conditions for *X* (SHR activator) and *Y* (SHR repressor), respectively (Supplementary Table 8).

For the production rate of SHR dimer in the CEI and QC (*k*_*2e*_ and *k*_*2q*_, respectively), the FPKM values of SCR were used to determine the value for *C*_*0*_, which is the concentration of SCR at which SHR dimer begins to form. It was assumed that the concentration of SCR required for SHR dimer formation is the same in both the CEI and QC. The value of SCR at 4D 16H (FPKM of 10.04) was taken as the steady state value because after this time SCR levels began to decrease. This value was then divided by 2 based on the assumption that *C*_*0*_, should be the same value in the CEI and QC (FPKM of 5.02 in each cell). Finally, this value was multiplied by 0.6 as previous results show that SHR dimer can form when SCR is at least 60% of its steady-state levels(16), resulting in the value *C*_*0*_=3. The steepness of the function, *s*, was chosen such that the production of SHR dimer reaches its maximum value as soon as SCR levels exceed *C*_*0*_ (as detailed in^23^). Thus, estimating *k*_*2e*_ and *k*_*2q*_ boiled down to estimating *L*_*e*_ and *L*_*q*_, which are the maximum values for *k*_*2e*_ and *k*_*2q*_ in the CEI and QC, respectively.

The parameters involved in SCR dimer formation in the CEI were set to 0 as the SCR dimer is at very low levels in the CEI (Supplementary Figure 1). Additionally, the diffusion coefficients of SHR (*a*_*e*_, *a*_*q*_) were not estimated as they were experimentally determined from RICS experiments in (19). The remaining sensitive parameters except for *k*_*1*_, *k*_*3e*_, and *k*_*3q*_ were set to constant values using the N&B data measured at 5D (Supplementary Table 3).

*k*_*1*_, *k*_*3e*,_ and *k*_*3q*_ were estimated from the time course of the root meristem using simulated annealing and Latin hypercube sampling as described in (27) This method produced 50 sets of initial parameter estimates, 33 of which did not converge, leaving 17 total estimated parameter sets (Supplementary Figure 11, Supplementary Table 1111). The average of these parameter values was used in the final model simulation (Supplementary Table 1111). MATLAB code used for simulated annealing is included in the data repository (see Data Availability).

Finally, the production terms for the SHR activator, SHR repressor, and SCR repressor (*k*_*9*_, *k*_*10*_, and *k*_*11e*_ and *k*_*11q*_ respectively) were set to a constant value at each time point based to minimize the model error compared to the time course data. Once SEU and WOX5 were identified as the SHR activator and repressor, *k*_*9*_ and *k*_*10*_ were re-estimated to minimize the error between the model and the time course expression data for these genes (Supplementary Table 10)

### Plant lines used in this study

The *wox5-1, seu-3, stk01, bzip17*, and SCRi lines are previously described in (16, 23, 43, 44). The 35S:GFP; UBQ10:mCherry; SHR:SHR-GFP; SCR:SCR-GFP; SHR:SHR-GFP, SCR:SCR:mCherry; and TMO5:3xGFP lines are described in (19). The WOX5:WOX5-GFP line is described in (30), the pCYCD6:GUS-GFP line is described in (7), and the SEU:SEU-GFP line is described in (45). The WOX5 TPT line (TPT_3.11260.1E) was obtained from the Arabidopsis Biological Resource Center (ABRC: https://abrc.osu.edu).

The SHR:SHR-GFP in *scr4* line was generated by crossing *scr4* mutant with SHR:SHR-GFP line. F2 lines homozygous for *scr4* mutation were selected by PCR using the oligos 5’-TTATCCATTCCTCAACTTCAGT-3’ and 5’-TGGTGCATCGGTAGAAGAAT-3’ which amplify a 302 DNA base pair fragment for the wild type and the oligos 5’-CTTATCCATTCCTCAACTCTATT-3’ and 5’-TGGTGCATCGGTAGAAGAATT-3’ which amplify a 296 DNA base pair fragment for the scr4 insertion. Selected lines were checked in the F3 generation for no segregation of SHR:SHR-GFP.

### Growth conditions

Seeds used for confocal microscopy were surface sterilized using fumes produced by a solution of 50% bleach and 1M hydrochloric acid, imbibed and stratified for 2 days at 4°C. After 2 days, the stratified seeds were plated and grown vertically at 22°C in long-day conditions (16-hrs light/8-hrs dark) on 1X Murashige and Skoog (MS) medium supplemented with sucrose (1% total). All plant roots were imaged at 5 days after plating unless otherwise noted. For the brassinolide (BL) treated plants, Columbia-0 (Col-0), pSHR:SHR:GFP, and pSCR:SCR:GFP seeds were prepared as described above but were plated on ½ MS medium plates with no sucrose and 4 nM Brassinolide. For RNAseq experiments, seeds were wet sterilized using 50% bleach, 100% ethanol, and water. Seeds were imbibed and stratified for 2 days at 4°C. After 2 days, the stratified seeds were plated on Nitex mesh squares on top of 1X MS medium with 1% sucrose. Seeds were plated and grown vertically at 22°C in long-day conditions.

### Confocal microscopy

Confocal microscopy was completed using a Zeiss LSM 710. Both 488nm and 570nm lasers were used for green and red channel acquisition, respectively. A solution of 10μM propidium iodide was used to stain cell walls for visualization. mPS-PI staining to visualize starch granules was performed as described in (46). For the N&B acquisition, 12-bit raster scans of a 256×256 pixel region of interest were acquired with a pixel size of 100nm and a pixel dwell time of 12.61μs as described in (47). For pCF acquisition, 8-bit line scans of a 32×1 pixel region of interest were acquired with a pixel size range of 40-100nm and a pixel dwell time of 12.61us as described in (47). Heptane glue was used during N&B and pCF acquisition to prevent movement of the sample as described in (47, 48).

### Number and Brightness (N&B) analysis

Analysis of the raster scans acquired for N&B was performed using the SimFCS software (47, 49) (https://www.lfd.uci.edu/globals/). The 35S:GFP and UBQ10:mCherry lines were used to calibrate software parameters. We used a region of interest of 64×64 or 128×128 pixels to analyze the CEI or QC cells, respectively. The S-factor for GFP (2.85) and for mCherry (0.86) were calculated first to normalize the autofluorescence/background region of the images (Supplementary Figure 8). After the S-factor was set, the monomer brightness of GFP (0.32) and mCherry (0.26) were measured and found to be similar to what was previously measured in the Arabidopsis root (19) (Supplementary Figure 8). To determine the possible stoichiometries of the SHR-SCR complex, a binding score was determined based on the colors of the stoichiometry histogram (Supplementary Figure 2) where 6 (red) corresponds to the highest amount of binding and 0 (blue) corresponds to no binding.

### RNAseq on mutant lines

For the *wox5-1* and *seu-3* RNAseq experiments, approximately 5mm of the root tip was collected. RNA was extracted using the RNeasy Micro Kit (Qiagen). cDNA synthesis and amplification were performed using the NEBNext Ultra II RNA Library Prep Kit for Illumina. Libraries were sequenced on an Illumina HiSeq 2500 with 100 bp single-end reads. Reads were filtered and mapped as described in the previous section. Differential expression was calculated using CuffDiff (50) with a cutoff of q<0.05 and fold change >2. Raw reads and FPKMs are available from GEO: https://www.ncbi.nlm.nih.gov/geo/query/acc.cgi?acc=GSE104945.

### qPCR

qPCR on 5 day old Col-0, *seu-3*, and *wox5-1* root tips was performed as described in (27). For each mutant, qPCR was performed on two technical replicates of two independent RNA samples (biological replicates). Results were comparable across biological replicates. Gene-specific primers used are provided in Supplementary Table 12.

### Sign score calculation

The sign score was used to compare the time course expression of candidate genes with the mathematical model prediction. The time course expression for the predicted SHR activators and repressors was obtained from the RNAseq time course of the root meristem. Replicate 1 was used for all of the candidate genes as only this replicate shows coexpression of SHR and SCR over time, which is supported by experimental evidence that SHR and SCR activate SCR expression. The sign score of the gene candidates compared to the model prediction was then calculated. The sign score is defined as the change in the model prediction between two time points (+1 for positive change, 0 for zero change, and −1 for negative change) times the change in the gene time course expression between the same two time points. Thus, the genes with a temporal pattern that is exactly the same as the model prediction will have the maximum possible sign score of 5. A threshold of 1.1 was used to determine a change in expression, meaning that gene expression must change at least 1.1-fold between time points in order to be recorded as a positive or negative change.

### Transcription Factor Yeast Two-Hybrid Assay

The Yeast Two-Hybrid assay was performed as described for the Yeast One-Hybrid assay in (23) with the following modifications. The SEU ORF was cloned into pGBKT7 (Clontech) and transformed into yeast strain AH109. SEU AH109 (bait) and AH109 empty vector were mated to the TF library described in (23) (Supplementary Table 6) and selected on SD-LeuUra. Four independent successful matings were replica-plated to quadruple dropout SD-LeuUraHisAde with and without 100 µg/mL 3-amino-1,2,4-triazol (3AT) and scored for colony growth over a period of four days. Positive interactions were scored as those matings with colony growth on SEU but not empty vector control plates.

### Corrected Total Cell Fluorescence (CTCF) Measurements

Corrected Total Cell Fluorescence (CTCF) measurements were calculated as described in (27). When measuring the CTCF for plants expressing pCYCD6:GUS:GFP, only CEI and CEI daughter cells were used for analysis. When calculating the CTCF of pCYCD6:GUS:GFP expression in *seu-3* and *wox5-1* mutants, there was a wide range in the measured CTCFs. Thus, to compare the samples, CTCF values were separated into three groups, “low” (CTCF/area < 1), “medium” (1 ≤ CTCF/area ≤ 5) and “high” values (CTCF/area > 5). The distributions of CTCF values between the samples were then compared using the Chi-Squared Test with Likelihood Ratio.

### Pair Correlation Function (pCF) analysis

Analysis of the line scans acquired for pCF was performed using the SimFCS software (47, 49) (https://www.lfd.uci.edu/globals/). Three technical replicates (pixel distance = 5, 7, and 9) were analyzed for each image as the cell walls are irregular in size, so changing the pixel distance can result in a different pCF carpet. For each technical replicate, a Movement Index (MI) was assigned based on if movement was detected in the carpet (arch pattern, MI=1) or not (no arch pattern, MI=0) as described in (19, 47). The technical replicates were then averaged for each biological replicate. The WOX5:WOX5:GFP images were analyzed separately in both directions. The 35S:GFP and TMO5:3xGFP control lines were analyzed only in the forward (left to right) direction as it has been previously shown that their movement index does not vary within the stem cell niche (19).

### Statistical analysis

All statistical analysis was performed using JMP software (https://www.jmp.com/). The Chi-Squared test with Likelihood ratio was used to determine if there was a significant difference in the distribution of CTCF values of CYCD6:GUS:GFP expression in *seu-3* and *wox5-1* mutants. For the remainder of the experiments, a Shapiro-Wilk Goodness of Fit test was used to determine normality of the data, and at least one sample in each experiment had a p<0.05 for this test, suggesting that not all of the data met normality assumptions. Therefore, the nonparametric equivalents of statistical tests (Wilcoxon test for 2 groups and Wilcoxon test with Steel-Dwass for greater than 2 groups) were used for all remaining statistical comparisons. Two-tailed comparisons were used, and p<0.05 was used for significance for all tests. The statistical test used for each experiment is reported in the figure legends. Exact sample sizes are reported in the figure legends and supplementary information. Exact p-values are reported in the supplementary information and in the data repository (see Data Availability).

## Data availability

All sequencing data are available on GEO at https://www.ncbi.nlm.nih.gov/geo/query/acc.cgi?acc=GSE104945 with access token ctwhoqsozdixlyb. All raw images and data, as well as all MATLAB code used for the model, are available at https://figshare.com/s/cc9c17b195b1d0894936

## Acknowledgments

We thank Erin Sparks, Cara Winter, and Philip Benfey for their insightful comments on this manuscript. Images in this manuscript were generated using the instruments and services at the Cellular and Molecular Imaging Facility (CMIF) at NCSU. Next-generation sequencing was performed by the Genomic Sciences Laboratory (GSL) at NCSU.

## Author Contributions

NMC and R. Sozzani conceptualized the study and designed the experiments. NMC and APF performed transcriptional profiling. NMC, LVdB, and ECN performed Scanning FCS. LVdB, ECN, and TTN performed qPCR and confocal imaging. APF and SGZ performed Y2H on SEU. NMC constructed and analyzed the mathematical model. BB and R. Simon contributed the WOX5:WOX5-GFP translational fusion. EBA and MMR contributed the SHR:SHR-GFP in *scr4* line. KLG provided the extensive background information on SHR movement and helped interpret experimental results. NMC made all main and supplementary figures. NMC and R. Sozzani wrote the paper, and all co-authors edited the paper.

## Supplementary Figures

**Supplementary Figure 1:**
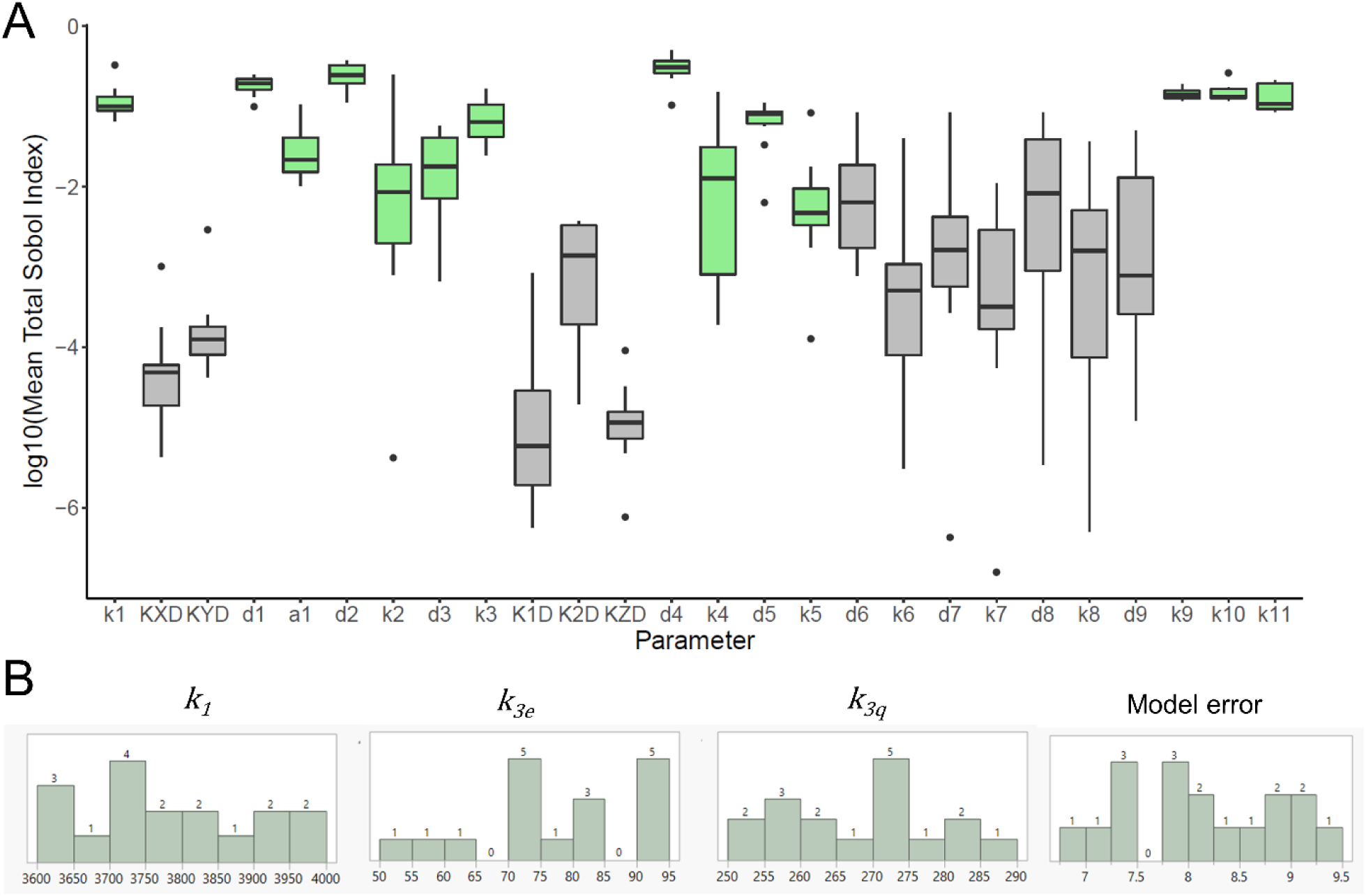
Sensitivity analysis and parameter estimation for SHR-SCR model. (A) Sensitivity analysis of SHR-SCR model. The Total Sobol Index was calculated for 10 technical replicates. The y-axis is displayed on a log10 scale. Green parameters are significantly sensitivity compared to *k*_*6*_, p<0.05, Wilcoxon with Steel Dwass. (B) Parameter estimation of SHR monomer production (*k*_*1*_) and SCR monomer production (*k*_*3e*,_ *k*_*3q*_) from time course dataset. The distribution of 17 best-fitting parameter sets is shown along with the model error for each set (far right histogram). Numbers above each bar represent the number of parameter sets in that range.

**Supplementary Figure 2:**
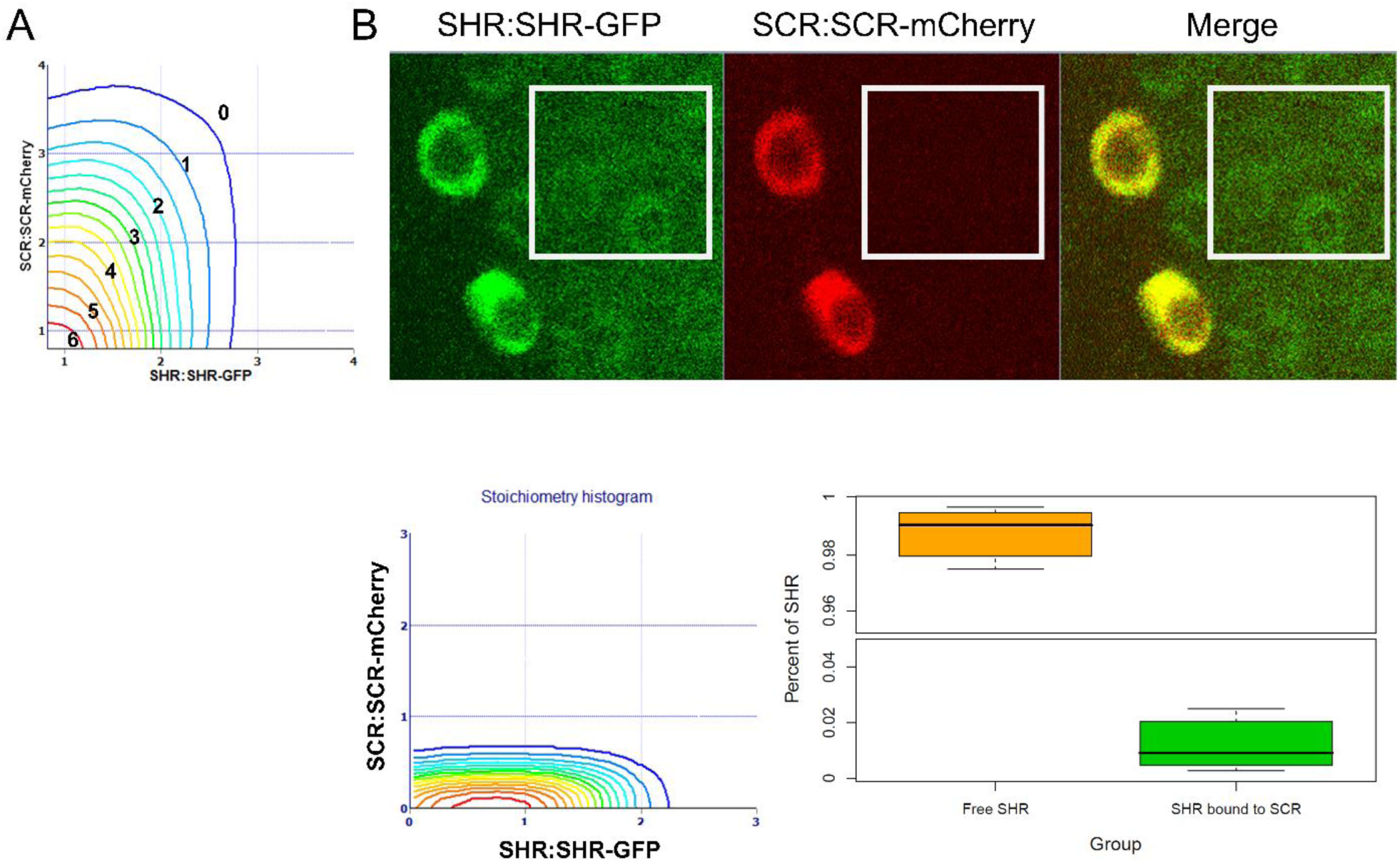
Cross N&B analysis in vasculature. (A) Quantification of binding score from Cross N&B output. Numbers correspond to the binding score given to complex stoichiometries depending on the color of the line in the stoichiometry plot. For example, red lines correspond to a score of 6, orange lines to 5, and so on. (B) (top) Representative SHR:SHR-GFP (left), SCR:SCR-mCherry (middle), and merged (right) image in 5 day old root. White box represents region of interested used for Cross N&B analysis. (bottom) (left) Stoichiometry histogram for this representative image. (right) Quantification of “free SHR” (not bound to SCR, orange) and SHR bound to SCR (green) in n=5 roots.

**Supplementary Figure 3:**
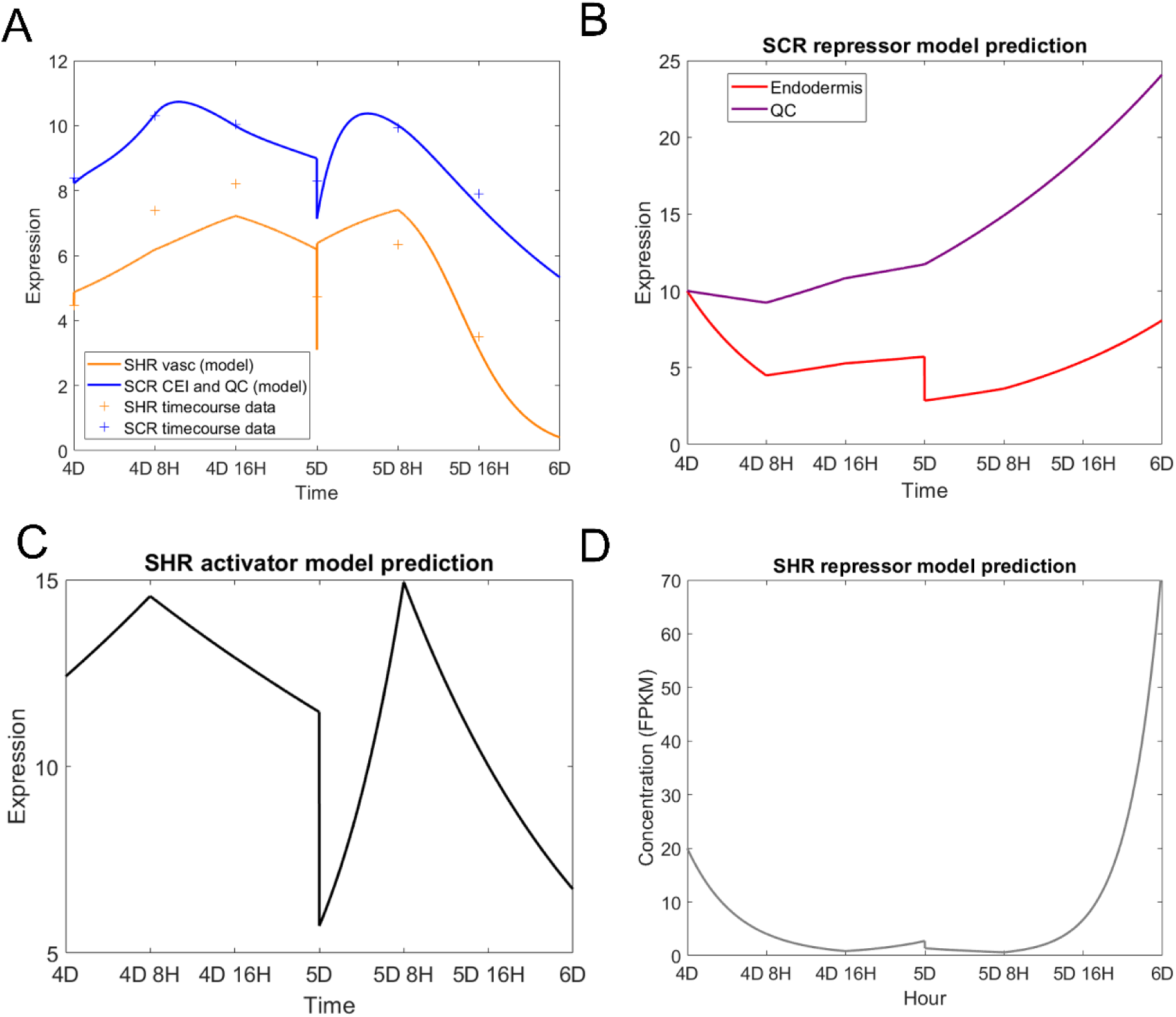
Model prediction of SHR, SCR, and their upstream SHR and their upstream regulators. (A) Model prediction of SHR in the vasculature (orange, solid line) and SCR in the CEI and QC (blue, solid line). Daggers represent experimental data used to fit model. (B) Model prediction of the SCR repressor in the endodermis (red) and QC (purple). (C,D) Model prediction of the SHR repressor (C) and activator (D) in the vasculature.

**Supplementary Figure 4:**
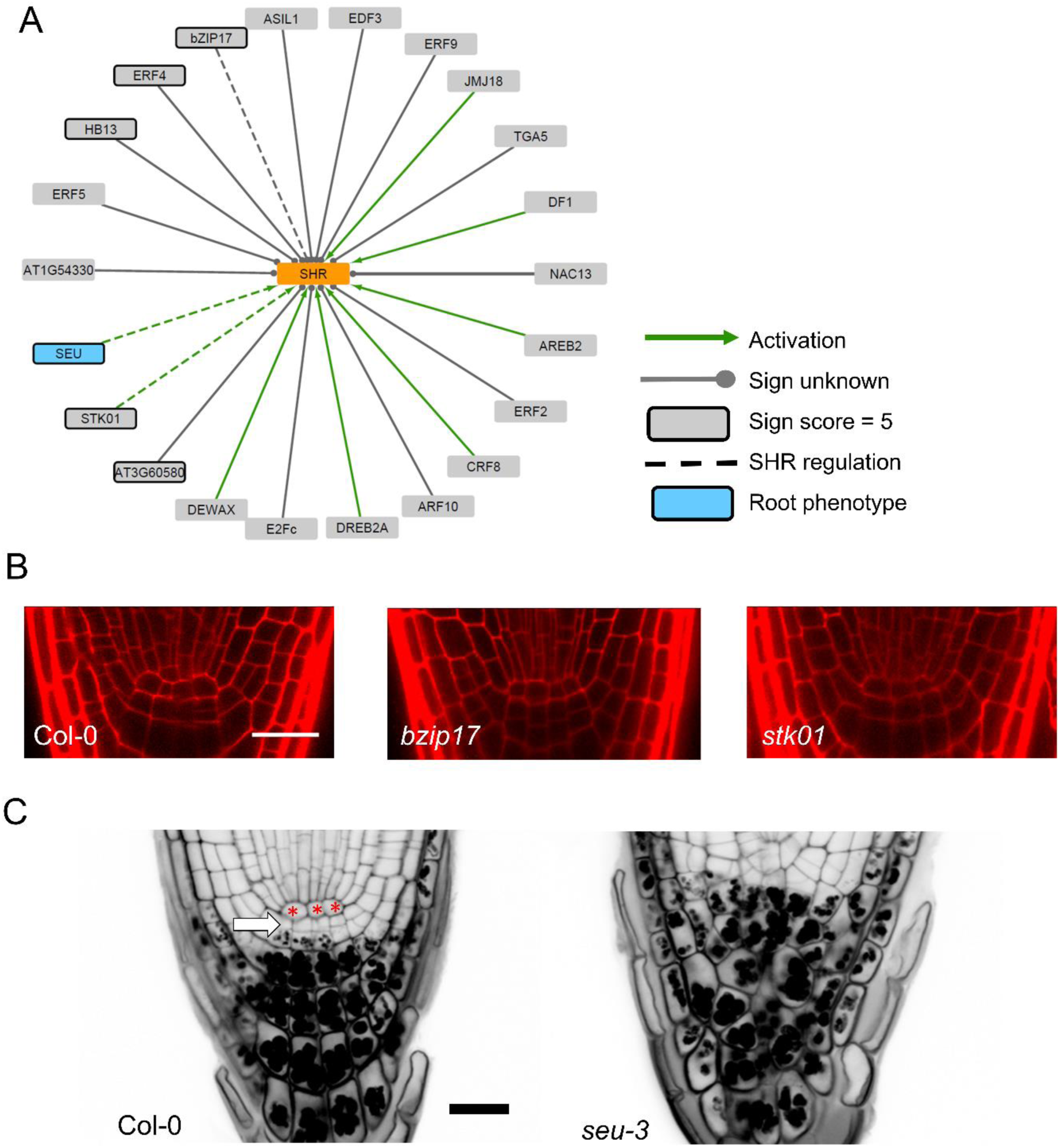
Putative activators of SHR. (A) Network of tested putative SHR activators. Black solid boxes indicate the 6 genes with the maximum possible sign score of 5 (black borders). Dashed lines indicate three out of the 6 genes showing low SHR levels in their mutant lines. Blue box indicates one of the 3 genes, SEU, whose mutant shows root stem cell disorganization. (B) Confocal images of putative SHR activator mutants of interest. Cell walls are stained with propidium iodide (PI). Scale bar = 20 µm. (C) mPSPI staining of *seu-3* mutants. Red * denote QC cells. White arrow denotes columella stem cells. Scale bar = 20 µm.

**Supplementary Figure 5:**
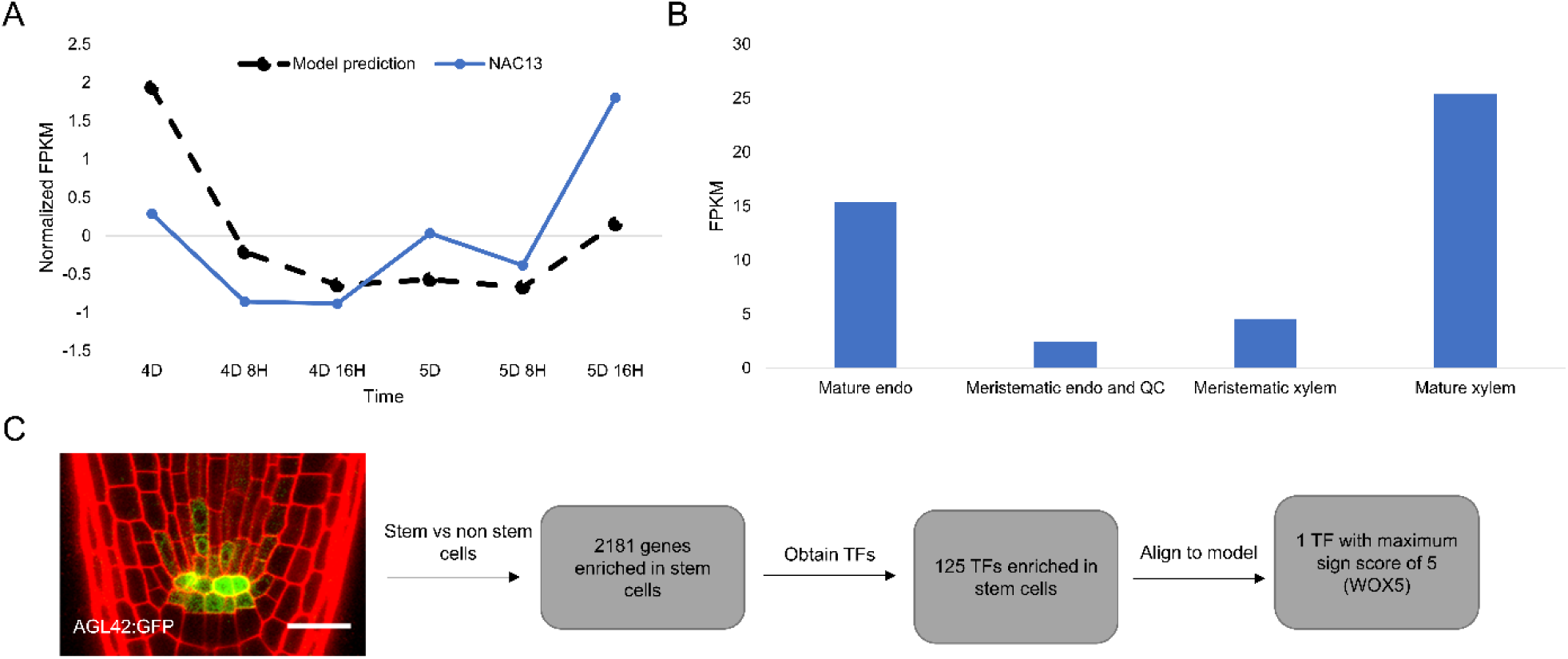
Identifying putative SHR repressors. (A) Comparison of model prediction for repressor (black, dashed line) and NAC13 normalized FPKM from the time course dataset (blue, solid line). (B) Average FPKM of NAC13 in different regions of the root meristem. (C) Workflow of obtaining a transcriptomic profile of the root stem cells. GFP positive (stem cells) and negative (non-stem cells) cells were collected from roots expressing AGL42:GFP. Cuffdiff identified 2181 genes (125 TFs) enriched in the stem cells. The expression of the 125 TFs was aligned to the model prediction of the SHR repressor and identified 1 TF with the highest sign score, WOX5.

**Supplementary Figure 6:**
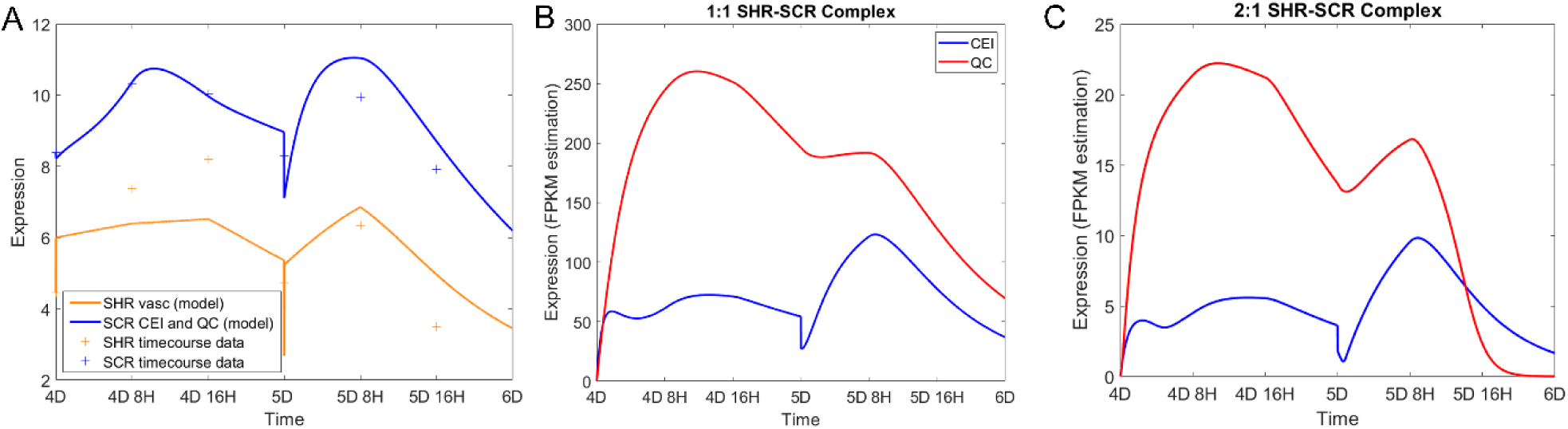
Final model simulation incorporating parameters estimated from SEU and WOX5 time course data. (A) Model prediction of SHR and SCR monomer. Solid lines represent model simulation, and daggers represent experimental data. (B,C) Model prediction of 1 SHR: 1 SCR (B) and 2 SHR: 1 SCR (C) complex in the QC (red) and CEI (blue).

**Supplementary Figure 7:**
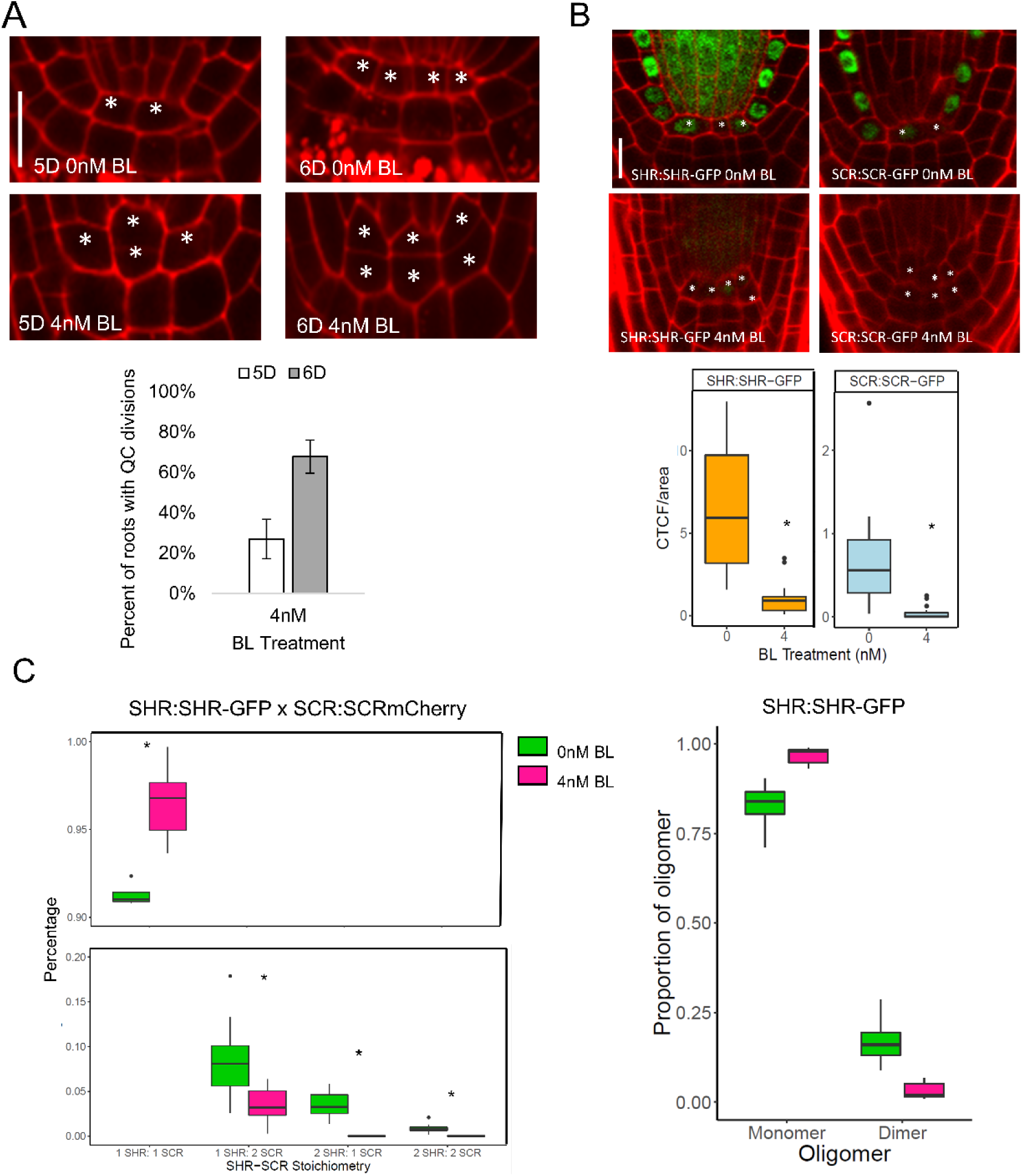
Effect of BL treatment on QC division, SHR and SCR expression, and SHR-SCR complex stoichiometry. (A) (top) Representative images of 5D (left half) and 6D (right half) plants with or without 4nM BL. * denotes QC cells. Scale bar = 10 µm. (bottom) Quantification of QC divisions in 4 nm BL treated roots at 5D (white bar) and 6D (gray bar). (B) (top) Representative images of 5 day old SHR:SHR-GFP (left half) or SCR:SCR-GFP (right half) with or without 4nM BL. * denotes QC cells. Scale bar = 10 µm. (bottom) Corrected total cell fluorescence (CTCF) of SHR:SHR-GFP (n=18) and SCR:SCR-GFP (n=16) in 0 and 4 nM BL treated plants. Black dots represent outliers. * denotes p < 0.05, Wilcoxon Test. (C) Proportion of SHR-SCR stoichiometries (left) and SHR oligomers (right) in the QC of plants grown in 0nM (green) and 4nM (pink) BL. * denotes p < 0.05, Wilcoxon test.

**Supplementary Figure 8:**
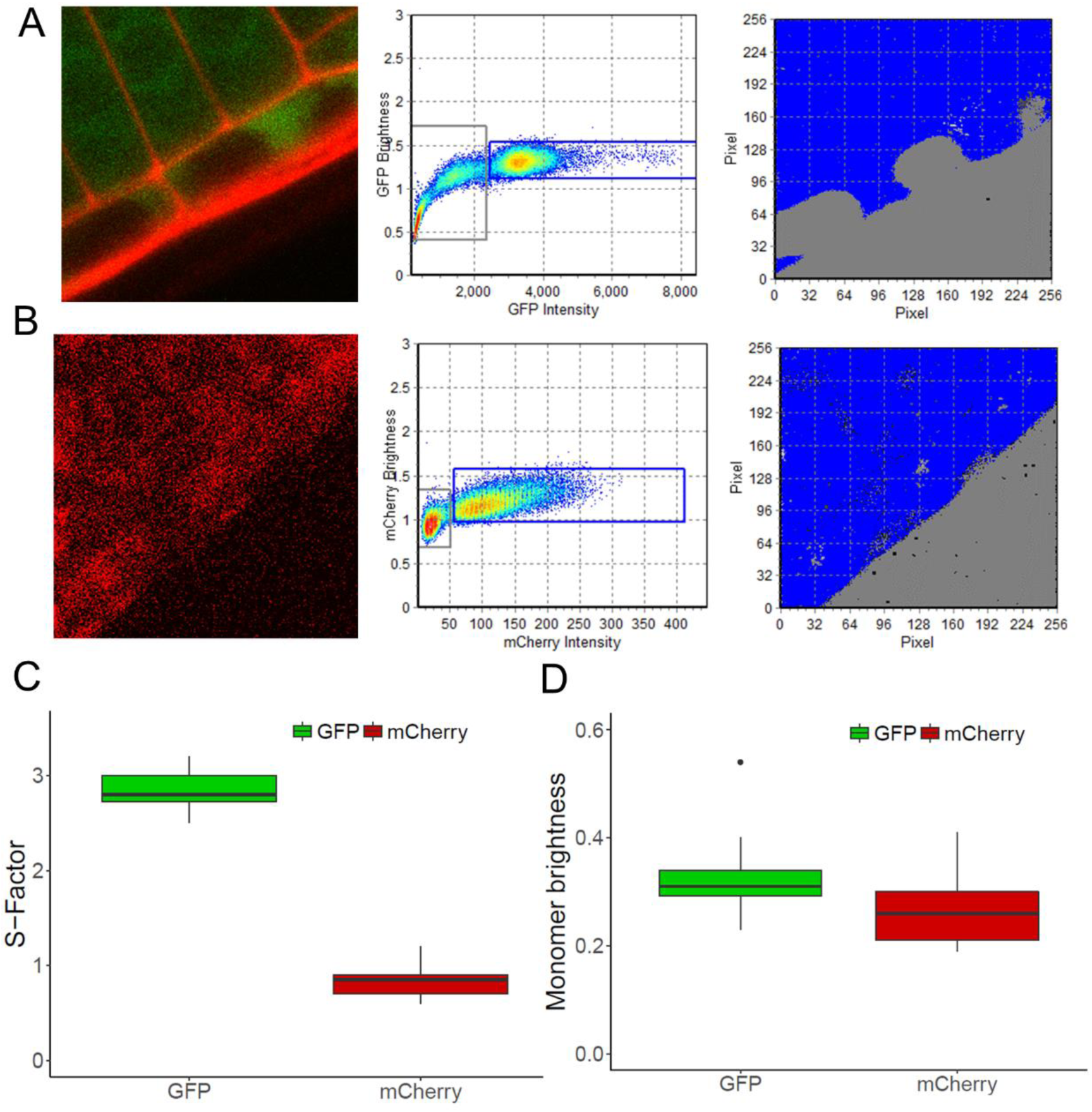
Calibration for N&B using free GFP and mCherry lines. (A, B) Representative image (left), brightness plot (middle), and false-colored image (right) for 35S:GFP (A) and UBQ10:mCherry (B) lines. Blue represents monomer, grey represents background. (C) S-factor calculated for 35S:GFP (green) and UBQ10:mCherry (red). (D) Monomer brightness calculated for GFP (green) and mCherry (red).

## Supplementary Tables

Supplementary Table 1: Sensitivity analysis of SHR-SCR model

Supplementary Table 2: Parameter values used in SHR-SCR model

Supplementary Table 3: FPKM values of SHR and SCR from the stem cell time course

Supplementary Table 4: Candidate SHR regulators and their sign scores

Supplementary Table 5: Differentially expressed genes in the *seu-3* mutant (uploaded as separate Excel file due to size)

Supplementary Table 6: Sign scores of the 125 TFs differentially expressed in the stem cells

Supplementary Table 7: Differentially expressed genes in the *wox5-1* mutant (uploaded as separate Excel file due to size)

Supplementary Table 8: SHR-SCR model parameters estimated using SEU and WOX5 time course data.

Supplementary Table 9: Yeast Two-Hybrid results (uploaded as separate Excel file due to size)

Supplementary Table 10: Expression of SCR in the QC and endodermis

Supplementary Table 11: Parameter sets generated from time course estimation.

Supplementary Table 12: Primers used in this study.

